# Rit2 silencing in dopamine neurons drives a Parkinsonian phenotype

**DOI:** 10.1101/2023.04.26.538430

**Authors:** Patrick J. Kearney, Yuanxi Zhang, Yanglan Tan, Elizabeth Kahuno, Tucker L. Conklin, Rita R. Fagan, Rebecca G. Pavchinskiy, Scott A. Shafer, Zhenyu Yue, Haley E. Melikian

## Abstract

Parkinson’s disease (PD) is the second most prevalent neurodegenerative disease and arises from dopamine (DA) neuron death selectively in the substantia nigra pars compacta (SNc). Rit2 is a reported PD risk allele, and recent single cell transcriptomic studies identified a major RIT2 cluster in PD DA neurons, potentially linking Rit2 expression loss to a PD patient cohort. However, it is still unknown whether Rit2 loss itself is causative for PD or PD-like symptoms. Here we report that conditional Rit2 silencing in mouse DA neurons drove motor dysfunction that occurred earlier in males than females and was rescued at early stages by either inhibiting the DA transporter (DAT) or with L-DOPA treatment. Motor dysfunction was accompanied by decreased DA release, striatal DA content, phenotypic DAergic markers, DA neurons, and DAergic terminals, with increased pSer129-alpha synuclein and pSer935-LRRK2 expression. These results provide the first evidence that Rit2 loss is causal for SNc cell death and a PD-like phenotype, and reveal key sex-specific differences in the response to Rit2 loss.

## Introduction

Parkinson’s disease (PD) is a complex, progressive, neurodegenerative disorder characterized by SNc DA neuron (DAN) death^1,2^. PD prevalence is higher in males and PD symptoms often go unnoticed until >75% of SNc neurons have died^3,4^. Phenotypically, PD patients exhibit profound motor impairment that includes bradykinesia, resting tremor, muscular rigidity, lack of coordination, and postural instability^5^. These symptoms are due to SNc DAN cell death and concomitant diminished striatal DA signaling, and PD therapeutic strategies typically aim to boost DA production in the remaining DAN population^6^.

Rit2 (AKA: Rin, **<u>R</u>**as-like **i**n **<u>n</u>**eurons) is a small, neuronal, ras-like GTPase with enriched expression in SNc DANs^7^. Rit2 directly interacts with the DA transporter (DAT), and is required for regulated DAT membrane trafficking^8–10^. In cell culture models, Rit2 is required for EGF- and NGF-mediated neurite outgrowth, NGF-mediated ERK phosphorylation, and cell viability^11–14^. Genome-wide association studies (GWAS) link Rit2 genetic anomalies to PD^15–26^, as well as to other neuropsychiatric disorders including essential tremor^24^, schizophrenia^24,27^, autism spectrum disorder^28,29^, bipolar disorder^24^, and speech delay^30^. Significant Rit2 mRNA decreases were reported in postmortem PD patient SNc^31^, and a recent scRNAseq study identified a major transcriptomic cluster in SNc DANs from PD subjects with striking Rit2 expression loss^32^. Moreover, Rit2 overexpression is sufficient to rescue cellular and behavioral deficits in an α- synuclein (α-syn) mouse PD model^33^. Together, these findings suggest that Rit2 plays a key role in DAN function and viability, and may be a critical factor in PD pathogenesis. However, it is still completely unknown whether Rit2 loss is, itself, causative for DAN degeneration and/or PD symptomology. Here, we leveraged our previously described approach for conditional Rit2 silencing in mouse DANs^8,9,34^ to directly test this possibility. Our results point to a direct role for Rit2 in DA neuron viability, and that Rit2 loss drives a PD-like phenotype with sex-specific differences.

## Results

### Rit2 silencing leads to sex-specific motor dysfunction

We previously leveraged the TET-off approach to conditionally silence Rit2 in Pitx3*^IRES-tTA^* midbrain DANs^8,9,34^, and reported that DAergic Rit2 silencing in both VTA and SNc for 4-6 weeks had no significant effect on either male or female baseline locomotion in the open field^34^. Given that PD is typically late onset, we aimed to comprehensively investigate the impact of conditional Rit2 knockdown (KD) in DANs on more complex motor behaviors at either short-term (4-6 weeks; ST) or long-term (5-6 months; LT) timepoints in both male and female mice. Pitx3*^IRES-tTA^* mouse VTA were bilaterally injected with AAV9-TRE-shRit2, which we previously reported drives shRit2 expression selectively in DANs in both VTA and SNc, due to AAV9 spread^9,34^. Consistent with our previous reports, AAV9-TRE-shRit2 significantly decreased Rit2 mRNA in both ST and LT male and female *Pitx3^IRES-tTA^* mouse midbrains, as compared to AAV9-TRE-eGFP injected controls (Supplemental Figure 1A-D). We previously reported that ST Rit2 KD had no effect on baseline locomotion. Similar to our previous finding in ST shRit2 mice, LT Rit2 KD likewise did not significantly affect horizontal locomotion in either male or female mice (Supplemental Figure 1E,F), nor was their fine movement significantly affected (Supplemental Figure 1G,I). There was additionally no change in male vertical motion (Supplemental Figure 1H), whereas female mice exhibited significantly increased vertical locomotion (Supplemental Figure 1J).

We next asked whether either ST or LT DAergic Rit2 KD impacted more complex motor behaviors, such as motor learning and coordination, assessed on rotarod and challenge balance beam assays. In male mice, both ST and LT DAergic Rit2 KD significantly decreased performance on the accelerating rotarod compared to controls (Figure 1A,B), whereas female mouse rotarod performance was not significantly affected by either ST or LT DAergic Rit2 KD (Figure 1C,D). Accelerating rotarod deficits could be due to either learning or coordination deficits. To discriminate between these possibilities, we assessed performance on the fixed- speed rotarod and challenge balance beam. Despite their accelerating rotarod deficits, male ST shRit2 mice did not exhibit any significant deficits on either the fixed-speed rotarod (Figure 1E), or challenge balance beam (Figure 1I,K), as compared to controls. Similar to males, female ST shRit2 mice did not exhibit any significant deficits on either the fixed-speed rotarod (Figure 1G) or challenge balance beam (Figure 1J,L), as compared to controls. We further assessed both mouse gait and grip strength following ST Rit2 KD, and found no differences between shRit2 mice and controls, in either males or females (Supplemental Figure 2).

**Figure 1.**
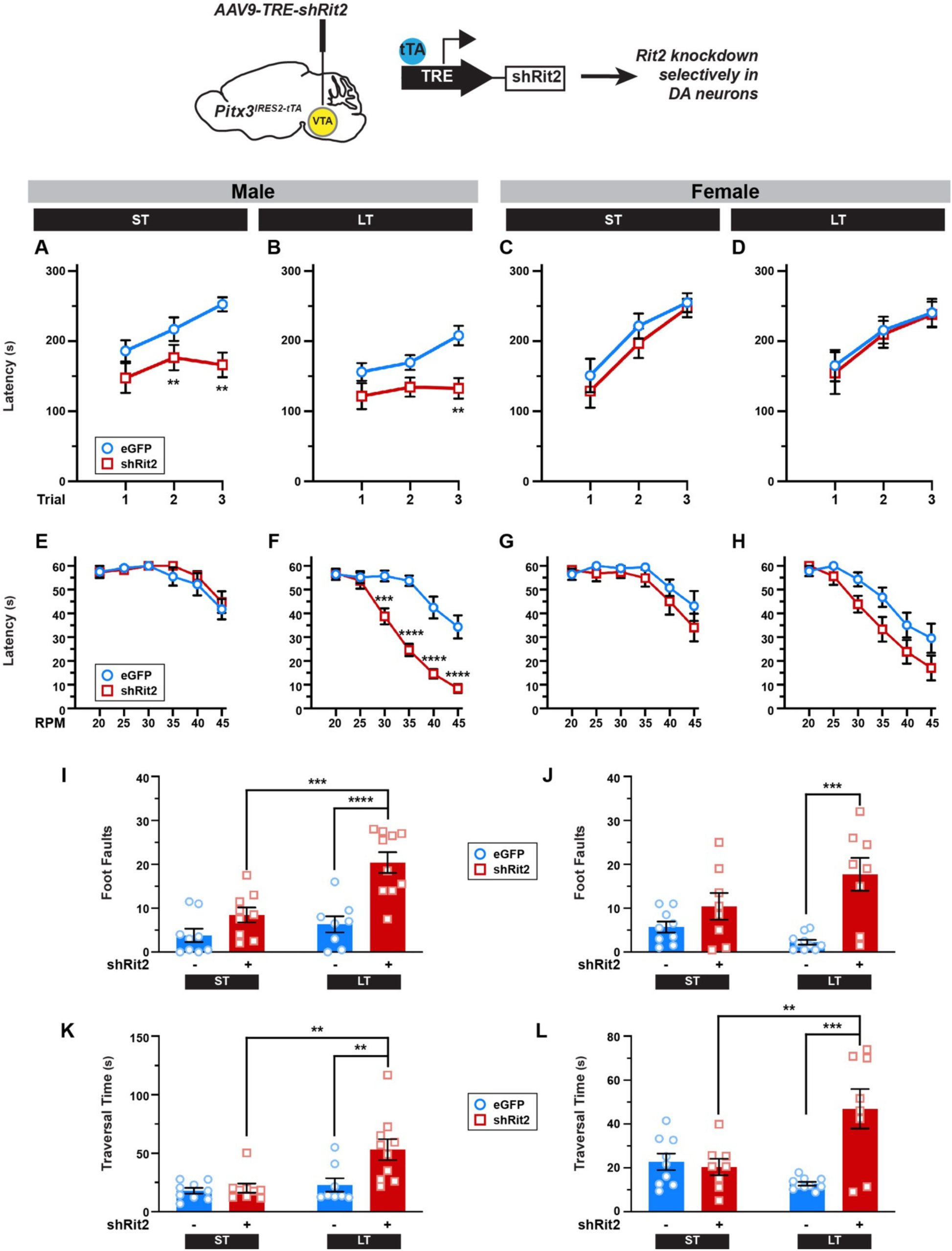
Conditional Rit2 silencing in DA neurons leads to progressive, but differential, motor dysfunction in males and females. *Pitx3^IRES-tTA^* mouse midbrains were bilaterally injected with either AAV9-TRE-eGFP or -shRit2 and mice were assessed either 4-5 weeks (ST) or 25 weeks (LT) post-injection. Data were analyzed by two-way repeat measures ANOVA with Sidak’s multiple comparison test (A-H), or two-way ANOVA with Tukey’s multiple comparison test (I-L)*. Top: Experimental schematic. Pitx3^IRES-tTA^* mouse midbrains were bilaterally injected with either AAV9-TRE-eGFP or -shRit2 and mice were assessed either 4-5 weeks (ST) or 25 weeks (LT) post-injection. **(A-D)** *Accelerating Rotarod:* Mice were assessed over 3 consecutive trials as described in *Methods.* Rit2 silencing in DA neurons abolished motor learning in male ST (**A.** Trial: p=0.001, F_(2, 41)_= 8.95; Virus: p=0.002, F_(1, 24)_ = 11.69; trial x virus: p=0.03; *p<0.005; n=9- 11) and LT (**B.** Trial: p=0.01, F_(2, 34)_=5.22; Virus: p=0.02, F_(1, 17)_=7.02; **p=0.005, n=8-11) mice, but had no effect on ST (**C.** Trial: p=0.002, F_(2, 48)_=6.82; Virus: p=0.72, F_(1, 48)_=0.12; n=11-12) or LT (**D.** Trial: p=0.002, F_(2, 48)_=6.82, Virus: p=0.72, F_(1,48)_=0.12, n=8-10) female motor learning. **(E-H)** *Fixed-speed Rotarod:* Mice were assessed over the indicated consecutive speeds as described in *Methods***. E,F.** Rit2 silencing did not affect ST males (**E.** Rate: p<0.0001, F_(5,70)_=11.59, Virus: p=0.48, F_(1,14)_=0.53, n=8), but significantly diminished LT male rotarod performance (**F.** Rate: p<0.0001, F_(5,102)_=46.88, Virus: p<0.0001, F_(1,102)_=98.31, Rate x Virus: p<0.0001, F_(5,102)_=5.34; ***p<0.001, ****p<0.0001, n=8-11) **G,H.** Rit2 silencing had no effect on female rotarod performance at the ST timepoint (**G.** RPM: p<0.0001, F_(2, 35)_=12.79, Virus: p=0.22, F_(1,15)_=1.637; n=8) but significantly diminished performance at the LT timepoint (**H.** RPM: p<0.0001, F_(5,108)_=6.29; Virus: p=0.006, F_(1,108)_=12.48; n=10). **I-L.** *Challenge balance beam:* Mice were assessed on the challenge balance beam as described in *Methods.* Mean foot fault numbers **(I,J)** and beam traversal times (seconds) **(K,L)** were analyzed. Rit2 silencing had no effect on ST male foot faults, but increased LT male foot faults (**I.** Time: p=0.0007, F_(1,32)_=14.23; Virus: p<0.0001, F_(1, 32)_=23.83; Time x virus: p=0.02, F_(1, 32)_=6.01; ***p=0.004, ****p<0.0001, n=8-10) and traversal times (**K.** Time: p=0.004, F_(1,32)_=9.52; Virus: p=0.01, F_(1, 32)_=7.02; Time x virus: p=0.03, F_(1, 32)_=5.17; **p<0.01, n=8-10). Rit2 silencing had no effect on ST female foot faults, but increased LT female foot faults (**J.** Time: p=0.42, F_(1,31)_=0.68; Virus: p=0.0002, F_(1, 31)_=18.59; Time x virus: p=0.03, F_(1, 31)_=5.36; ***p=0.0003, n=8-10) and traversal times (**L.** Time: p=0.10, F_(1,31)_=2.85; Virus: p=0.003, F_(1, 31)_=10.64; Time x virus: p=0.0008, F_(1, 31)_=13.95; **p=0.005, ***p=0.0001, n=8-10).

Given that ST Rit2 KD only affected male accelerating rotarod performance, we next asked whether longer Rit2 silencing would lead to motor dysfunction in females, and/or development of motor dysfunction on other motor tasks. Following LT Rit2 KD, males continued to exhibit significantly poorer performance than controls on the accelerating rotarod (Figure 1B), and females still exhibited no significant effect on accelerating rotarod performance (Figure 1D). However, LT Rit2 KD drove a significant deficit in fixed-rotarod performance in both males (Figure 1F) and females (Figure 1H), and both foot faults and beam traversal times were significantly increased in both male (Figure 1I,K) and female (Figure 1J,L) LT shRit2 mice on the balance beam, as compared to both control and ST Rit2 KD mice. Gait analysis revealed that multiple gait parameters were unaffected by ST Rit2 KD in both males (Supplemental Figure 2A,E,I,M) and females (Supplemental Figure 2C,G,K,O). However, there was a differential impact on gait in males and females following LT Rit2 KD. LT shRit2 males completed significantly fewer gait analysis trials (Supplemental Figure 2B) and had significantly narrower forelimb stride widths (Supplemental Figure 2J), whereas LT shRit2 females had significantly wider hindlimb stride widths (Supplemental Figure 2L). Despite the observed coordination and gait deficits, all LT shRit2 mice also had significantly increased four-limb grip strength (Supplemental Figure 2R,T). Taken together, the behavioral data suggests that DAergic Rit2 is specifically required for male motor learning and that prolonged Rit2 suppression leads to motor coordination and gait deficits in both males and females.

### Short-term DAergic Rit2 silencing blunts DA release in males

Motor learning deficits in males in response to ST DAergic Rit2 silencing could be due to altered DA release. To test this possibility, we leveraged fast-scan cyclic voltammetry (FSCV) to measure both DA release and clearance in *ex vivo* dorsal striatal slices (Figure 2A,B). Given the viability issues inherent to acute brain slices prepared from older animals, we limited our FSCV studies to ST shRit2- and control-injected males, in which we observed a motor learning deficit. We^9^ and others^35^ previously reported that DRD2 autoreceptors significantly blunt DA transient amplitudes and accelerate DA clearance. Indeed, in control mice DA transient amplitudes were significantly smaller when evoked in ACSF as compared to those evoked in the presence of L- 741,626 (25nM, Figure 2C), a DRD2-specific antagonist, as we previously reported^9^. In shRit2 mice, DA transient amplitudes recorded in ACSF were not significantly different transients from control mice (Figure 2C). However, unlike control DA transients, DA amplitudes recorded in the presence of L-741,626 were not significantly greater than those recorded in ACSF, and were significantly smaller than amplitudes recorded in L-741,626 from control mice. Moreover, we previously reported that ST Rit2 silencing in males decreased DAT surface levels by ∼50% in dorsal striatum (DS). Despite this reduction in DAT, Rit2 silencing did not significantly affect DA clearance times (Figure 2D), and we still observed DRD2-mediated enhancement of DA clearance in slices from both control and shRit2 mice, suggesting that DRD2 signaling was intact (Figure 2D). Taken together, these results suggest that males locomotor deficits following ST Rit2 silencing may be, in part, due aberrant DA signaling.

**Figure 2.**
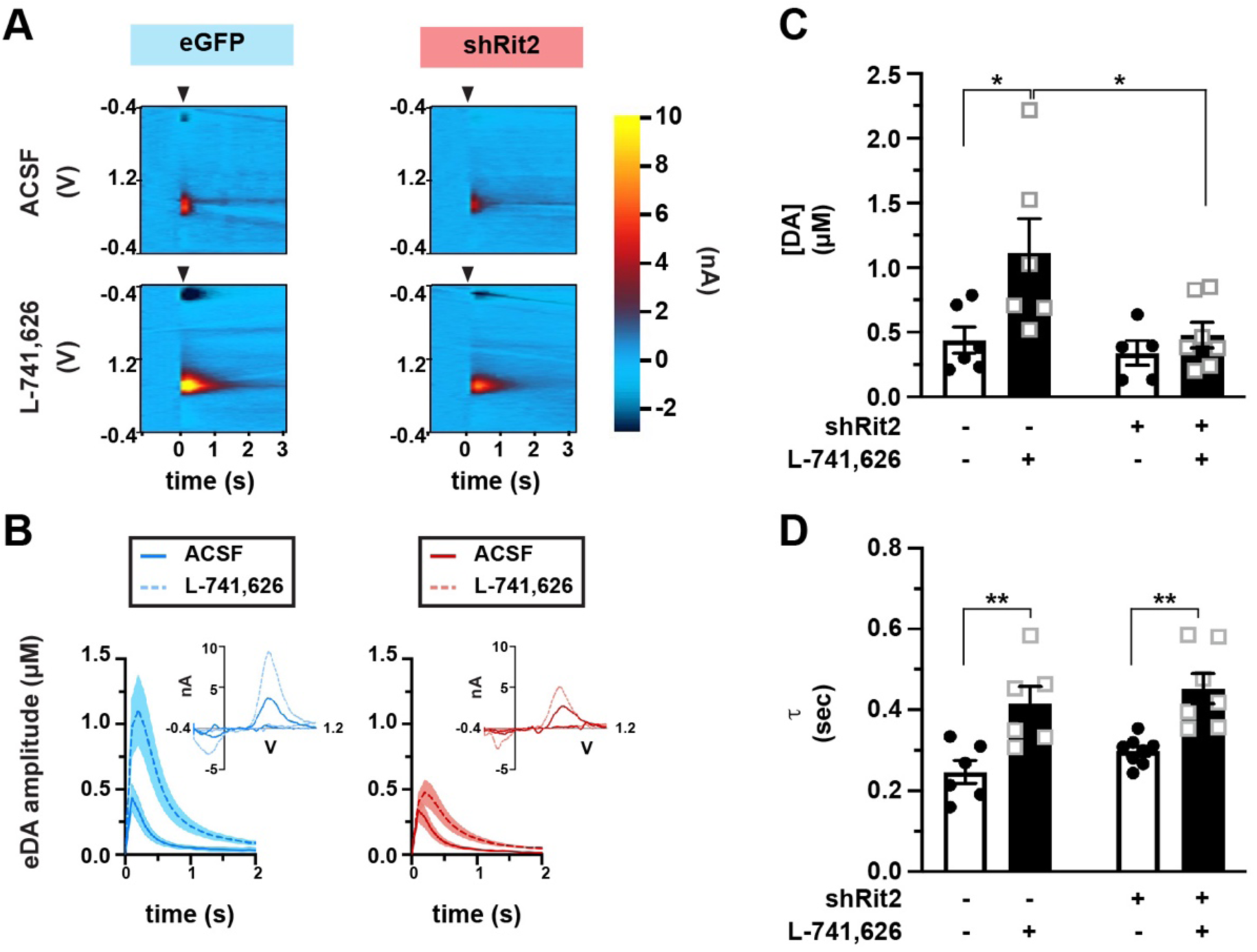
Short-term DAergic Rit2 silencing impacts DA release, but not clearance in males. *Ex vivo fast-scan cyclic voltammetry: Pitx3^IRES-tTA^* mouse VTA were bilaterally injected with either AAV9-TRE-eGFP (n=6) or AAV9-TRE-shRit2 (n=5-7) and electrically evoked DA transients were measured *ex vivo* in acute dorsal striatum as described in *Methods.* Values were analyzed by two-way ANOVA with Tukey’s multiple comparison test. Data were acquired from n=5-8 DS slices prepared from 3-4 independent mice per viral condition. **(A)** *Representative voltammograms:* Voltammograms displaying evoked current over voltage cycles and time, in slices from eGFP- and shRit2-injected mice, recorded in either ACSF or 25nM L-741,626, as indicated. Arrowheads indicate delivery of single, squared wave pulse. **(B)** *Dopamine transients:* Representative evoked DA transients in slices in slices from eGFP- and shRit2-injected mice, recorded in either ACSF or 25nM L-741,626, ±S.E.M. (shaded areas), as indicated. **(C)** *Average amplitudes:* DA transient amplitudes are presented in µM ±S.E.M. Virus: p=0.03, F_(1, 20)_=2.83; Drug: p=0.02, F_(1, 20)_=6.58; *p<0.05. **(D)** Average decay tau, presented in seconds ±S.E.M. Virus: p=0.16, F_(1, 23)_=2.08; Drug: p<0.0001, F_(1, 23)_=27.43; **p<0.01.

### Rit2 silencing suppresses the DAergic phenotype with earlier manifestation in males than females

Given our FSCV results, we hypothesized that motor deficits observed in ST males, and in both males and females following LT Rit2 silencing, could potentially be due to a loss in DAergic tone. To test this possibility, we first measured striatal DA content using mass spectroscopy in male and female dorsal (DS) and ventral (VS) striata following ST and LT Rit2 silencing. In ST shRit2 mice, total DA content was not significantly affected in either DS or VS from either male or female mice as compared to their respective controls (Figure 3A,C). However, DA content was significantly reduced in LT shRit2 male DS (Figure 3B) and LT shRit2 female VS and DS (Figure 3D) as compared to controls. Importantly, total striatal GABA content was not altered in male or female VS or DS at any timepoint (Figure 3E-H), demonstrating specific changes in DANs and not global changes in striatal neurotransmitter content.

**Figure 3.**
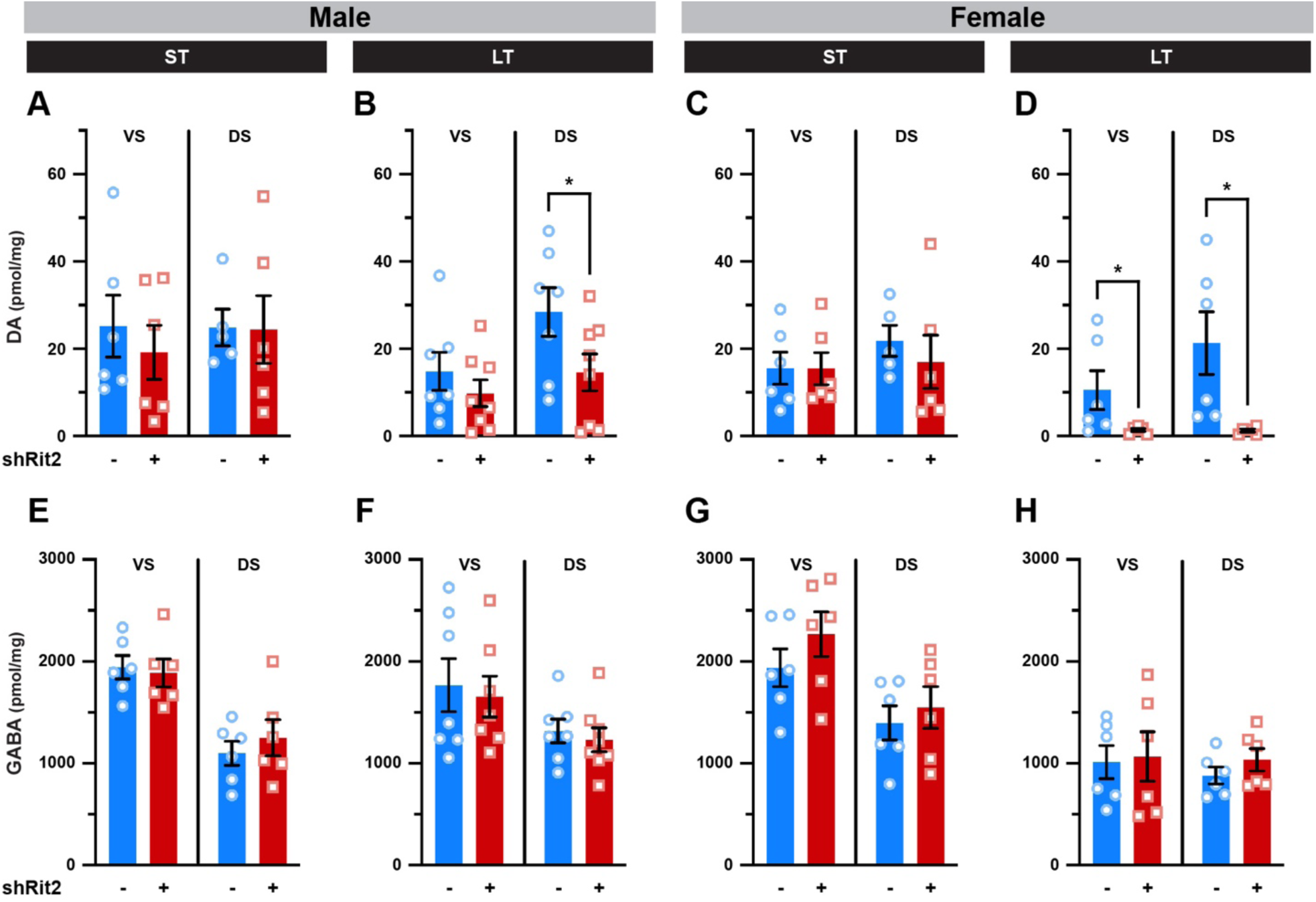
Long-term, but not short-term, Rit2 silencing decreases striatal DA content. Mass spectrometry. Dorsal (DS) and ventral (VS) striata were dissected from male and female control and shRit2 mice at the indicated timepoints, and total DA and GABA content were measured by LC/MS/MS as described in Methods. Data are presented as pmol of the indicated neurotransmitter per mg tissue, and were analyzed by unpaired, one-tailed (DA) or two-tailed (GABA) Student’s t test. N values indicate the number of independent striata. **(A-D)** *Striatal DA Content*: ST Rit2 silencing had no effect on DA content in male VS and DS (**A.** VS: p=0.27; DS: p=0.48, n=5-6), whereas LT Rit2 silencing decreased DA in DS, but not VS (**B.** VS: p=0.18; DS: p=0.03, n=7-8). ST Rit2 silencing had no effect on DA content in female VS and DS (**C.** VS: p=0.49; DS: p=0.26, n=5-6), but significantly decreased DA in LT females in both VS and DS (**D.** VS: *p=0.04; DS: *p=0.02, t test with Welch’s correction, n=6). **(E-H)** *Striatal GABA content.* Rit2 silencing had no effect on GABA content in either VS or DS in ST males (**E.** VS: p=0.75; DS: p=0.50, n=6), LT males (**F.** VS: p=0.74; DS: p=0.61, n=7-8), ST females (**G.** VS: p=0.28; DS: p=0.58, n=6), or LT females (**H.** VS: p=0.86; DS: p=0.29, n=6).

Given Rit2’s association with PD and the profound changes in motor function and DAergic tone observed with LT Rit2 silencing, we hypothesized that Rit2 silencing may decrease DAN viability. To test this possibility, we first measured DAergic gene and protein expression in isolated ventral midbrain (vMB) and striatum, respectively, following ST and LT Rit2 silencing. In males, RT- qPCR studies revealed that ST Rit2 KD significantly decreased tyrosine hydroxylase (TH) and DAT mRNA in vMB (Figure 4A,B), and quantitative immunoblotting revealed that striatal TH and DAT protein were also significantly reduced (Figure 4I,J). Unsurprisingly, following LT Rit2 KD, male TH and DAT vMB mRNA (Figure 4C,D) and striatal TH and DAT protein (Figure 4K,L) remained significantly diminished compared to controls. In females, TH and DAT mRNA (Figure 4E,F) and protein (Figure 4M,N) were unaffected following ST Rit2 KD. However, following LT Rit2 silencing females exhibited robust and significant loss in vMB TH and DAT mRNA (Figure 4G,H), as well as striatal TH and DAT protein (Figure 4O,P).

**Figure 4.**
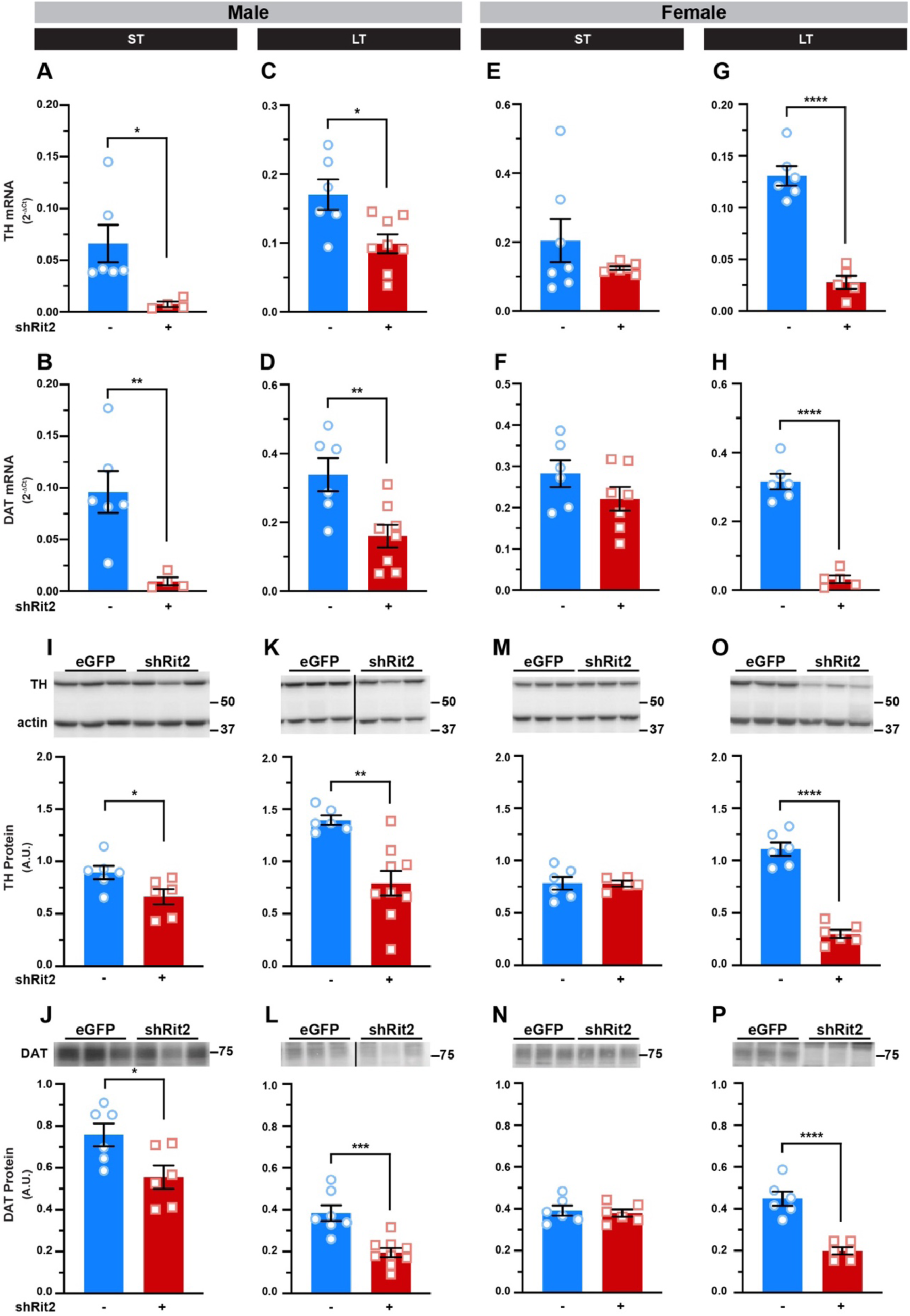
DAergic Rit2 silencing diminishes DAergic mRNA and protein expression. Ventral midbrain and striatum were dissected from male and female control and shRit2 mice at the indicated timepoints, and TH and DAT mRNA and protein were measured in ventral midbrain and striatal subregions, respectively, as described in Methods. Data were analyzed by unpaired, two-tailed Student’s t test. N values indicate the number of striata analyzed from independent mice (**A-D**) Male ventral midbrain RT-qPCR. TH (**A**. *p=0.02 with Welch’s correction, n=4-7) and DAT (**B**. **p=0.008 with Welch’s correction, n=4-6) mRNA were significantly decreased in ST shRit2 VM and both TH (**C.** *p=0.01, n=6-8) and DAT (**D.** **p=0.008, n=6-8) remained suppressed in LT shRit2 ventral midbrain as compared to eGFP controls. (**C-F)** Female ventral midbrain RT-qPCR. Neither TH (**E**. p=0.25 with Welch’s correction, n=7) nor DAT (**F**. p=0.19, n=6-7) mRNA were significantly affected in ST shRit2 ventral midbrain, whereas both TH (**C.** ****p<0.0001, n=5-6) and DAT (**D.** ****p<0.0001, n=5-6) were diminished in LT shRit2 ventral midbrain as compared to eGFP controls. (**I-L)** Male striatal protein. Top: Representative striatal immunoblots for each protein, showing 3 independent mouse lysates each for control and shRit2 mice. Molecular weight markers are indicated in kDa. TH (**I.** *p=0.04, n=6) and DAT (**J.** *p=0.03, n=6) protein levels were significantly decreased in ST shRit2 striata, and both TH (**K.** ***p=0.0008 with Welch’s correction, n=6-9) and DAT (**L.** ***p=0.0004, n=7-9) continued to be significantly decreased in LT shRit2 striata. (**M-P**) Female striatal protein. Top: Representative striatal immunoblots for each protein, showing 3 independent mouse lysates each for control (eGFP) and shRit2 mice. Neither TH (**M.** p=0.96) nor DAT (**N.** p=0.71, n=5-6) striatal protein levels were significantly diminished in shRit2 striata as compared to controls. In LT shRit2 striata, both TH (**O.** ****p<0.0001, n=6) and DAT (**P.** ****p<0.0001, n=6) protein levels were significantly reduced as compared to controls.

We further tested whether Rit2 silencing impacted TH activation, by measuring pSer40-TH via immunoblot. When normalized to actin, pSer40-TH was significantly reduced in ST shRit2 male striatum (Supplemental Figure 3A) and trended to decrease in LT shRit2 males (Supplemental Figure 3B). However, proportion of pSer40-TH to total TH was not significantly different in ST or LT shRit2 male mice as compared to controls (Supplemental Figure 3E,F), suggesting that functional TH regulation is intact, despite diminished total TH levels. In females, ST Rit2 silencing had no effect on pSer40-TH or the fraction of pSer40-TH (Supplemental Figure 3C,G). However, there was a drastic pSer40-TH loss following LT Rit2 silencing in females (Supplemental Figure 3D), as well as the fraction of pSer40-TH (Supplemental Figure 3H), suggesting that TH is dysregulated following LT Rit2 KD in females.

Given the profound losses in TH and DAT expression, as well as DA content, we further asked whether other characteristic ventral midbrain DAergic mRNAs were affected by shRit2. In both ST and LT shRit2 males, DRD2 and Pitx3 mRNA were significantly decreased (Supplemental Figure 4A,B,E,F), and Nurr1 was significantly diminished following LT, but not ST, Rit2 silencing (Supplemental Figure 4I,J). In females, ST Rit2 silencing did not significantly affect DRD2, Pitx3, or Nurr1 mRNA levels (Supplemental Figure 4C,G,K). However, by the LT timepoint all three DAergic markers were significantly diminished (Supplemental Figure 4D,H,L). Taken together these data demonstrate that DAergic Rit2 silencing results in downregulation of all DAergic genes, consistent with the loss in DAergic tone.

We additionally tested whether gene silencing in response to Rit2 KD was specific to DAergic genes or whether pan neuronal and/or ubiquitous genes are also affected by Rit2 silencing (Supplemental Figure 5). We measured vMB expression of the ubiquitously expressed Rit2 homolog, Rit1, and Vps35, a core retromer component that is also associated with PD. Surprisingly, both Rit1 and Vps35 gene expression were increased in ST, but not LT shRit2 males (Supplemental Figure 5A,B). Rit1 and Vps35 expression were unaffected in ST shRit2 females (Supplemental Figure 5C,G) but significantly increased at the LT timepoint (Supplemental Figure 5D,H).

### Prolonged DAergic Rit2 silencing increases PD markers in the striatum

Given Rit2’s association with PD, and the striking motor dysfunction we observed following Rit2 silencing, we next asked whether the loss in DAergic markers following Rit2 silencing was accompanied by an appearance of PD markers. To test this, we measured striatal αSyn and pSer129-αSyn levels, which markedly increase in idiopathic PD^36^. Total αSyn levels were not significantly affected in either ST or LT shRit2 males (Figure 6A,B) nor in LT females (Figure 6D), but were significantly increased in ST shRit2 females (Figure 6C). Importantly, shRit2 drove a significant increase in pSer129-αSyn in ST and LT shRit2 males (Figure 6E,F) and in LT females (Figure 6H), and strongly trended for an increase in ST females (Figure 6F). We additionally measured whether there were any changes in leucine-rich repeat kinase 2 (LRRK2), whose variants have been linked to PD and cause LRRK2 phosphorylation anomalies^37^. Total LRRK2 levels did not significantly differ from controls in all ST and LT shRit2 mice (Supplemental Figure 6). However, LT Rit2 KD significantly increased pSer935-LRRK2 in both LT males and females, while ST Rit2 KD did not significantly affect pSer935-LRRK2 in males or females. Together, these data indicate that in addition to profound motor deficits and DAN degeneration, LT DAergic Rit2 silencing drives an increase in both phosphorylated α-synuclein and LRRK2.

### Long term Rit2 silencing results in DAN degeneration

DAergic marker loss and pSyn increases may be due to cell death, or quiescence of the DAergic phenotype without cell death. To discriminate between these possibilities, we performed stereological counting to measure the number of total (Nissl stain) and dopaminergic (TH+) SNc neurons at the LT Rit2 KD timepoint, where both males and females exhibited significant motor dysfunction. In LT Rit2 KD males, there was no significant loss of total SNc neurons (Fig. 5A), and a strong trend towards decreased TH+ cells in the SNc (p=0.07, Fig. 5B). Moreover, when normalized to the total SNc cells, there was a significant decrease in the %TH+ cells in male SNc (Fig. 5C). In LT Rit2 KD females, there was a significant decrease in both total SNc neurons (Fig. 5D), as well as TH+ cells (Fig. 5E). There was, however, no difference in the % neurons that were TH+ in LT Rit2 KD females as compared to controls (Figure 5F), suggesting that the SNc neuronal losses occurred primarily in the DAergic population. Together, these results are consistent with DAN cell death following LT Rit2 KD.

**Figure 5.**
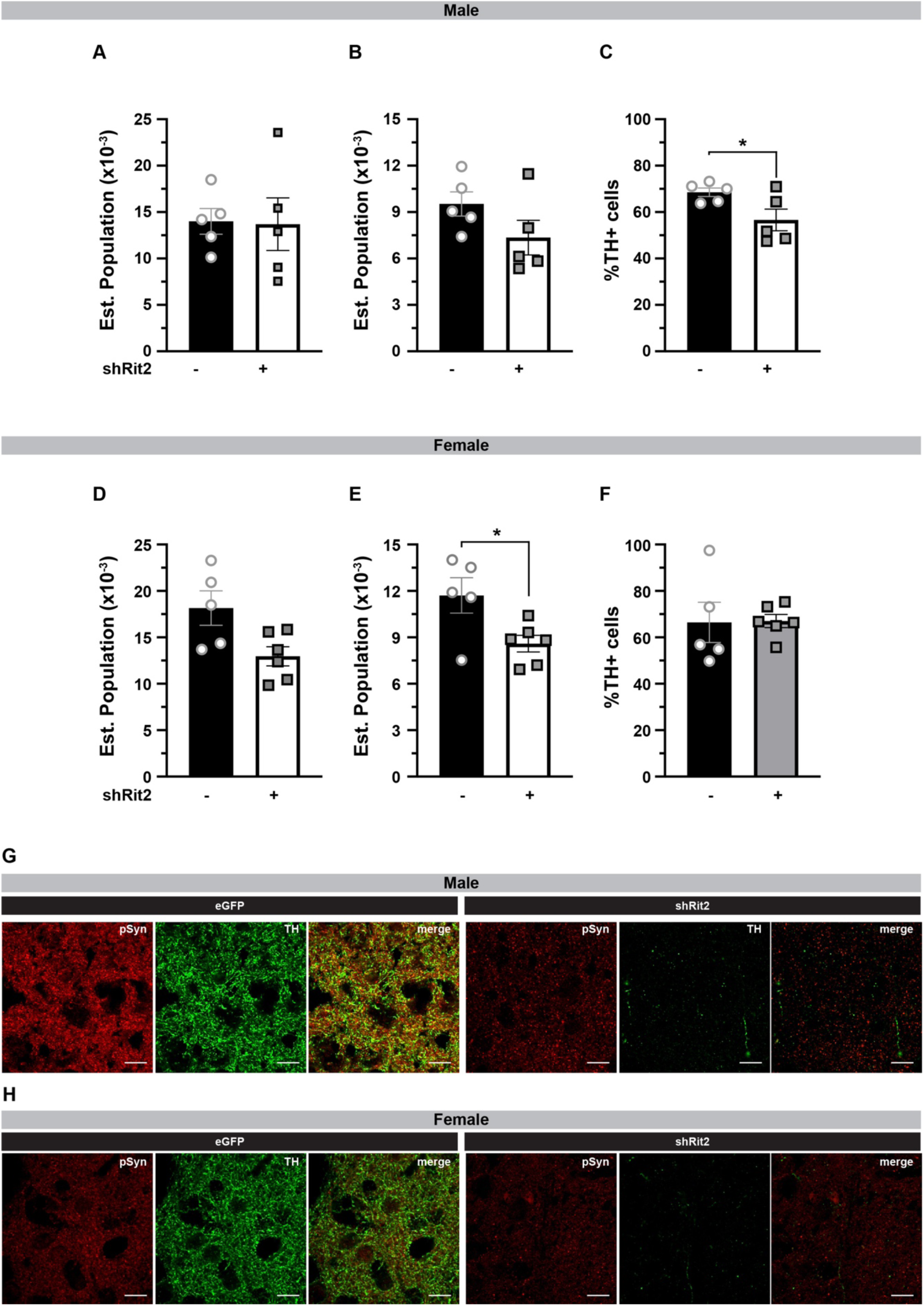
Prolonged DAergic Rit2 silencing decreases the substantia nigra DA neurons and striatal DAergic terminals. *Immunohistochemistry* Pitx3^IRES-tTA^ males and female mouse VTA were bilaterally injected with either AAV9-TRE-eGFP or AAV9-TRE- shRit2 and brains were fixed, sectioned, stained with the indicated dyes/antibodies and analyzed at the LT timepoint as described in *Methods.* **A-F.** *Stereological analysis of SNc neurons.* Midbrain sections were stained with cresyl violet and TH-specific antibodies. Total and TH+ cell numbers were counted as described in *Methods* and significance was tested with a two-tailed, unpaired, Student’s t test. N values indicate the number of independent mouse SNc analyzed. **A-C**. *Males:* Rit2 KD did not significantly decrease total Nissl+ neurons (**A.** p=0.46), trended towards a decrease in TH+ neurons (**B.** p=0.07) cells, and significantly decreased the %TH+ neurons (**C. \***p=0.02) in SNc, n=5. **D-F.** *Females:* Rit2 KD significantly decreased both total Nissl+ neurons (**D. \***p=0.02), and TH+ neurons (**E.** *p=0.01) cells, but did not significantly decrease the %TH+ neurons (**F.** p=0.48) in SNc, n=5-6. **G, H.** *Striatal immunohistochemistry.* Sections were co-stained for pSer129-Syn (red) and TH (green) and imaged by confocal microscopy as described in *Methods.* A single representative plane is presented from male (**G**) and female (**H**) dorsal striatum, from 2 independent mice for each virus and sex. Scale bars = 20µm. LT Rit2 KD dramatically decreased TH+ terminals in dorsal striatum in both males and females.

**Figure 6.**
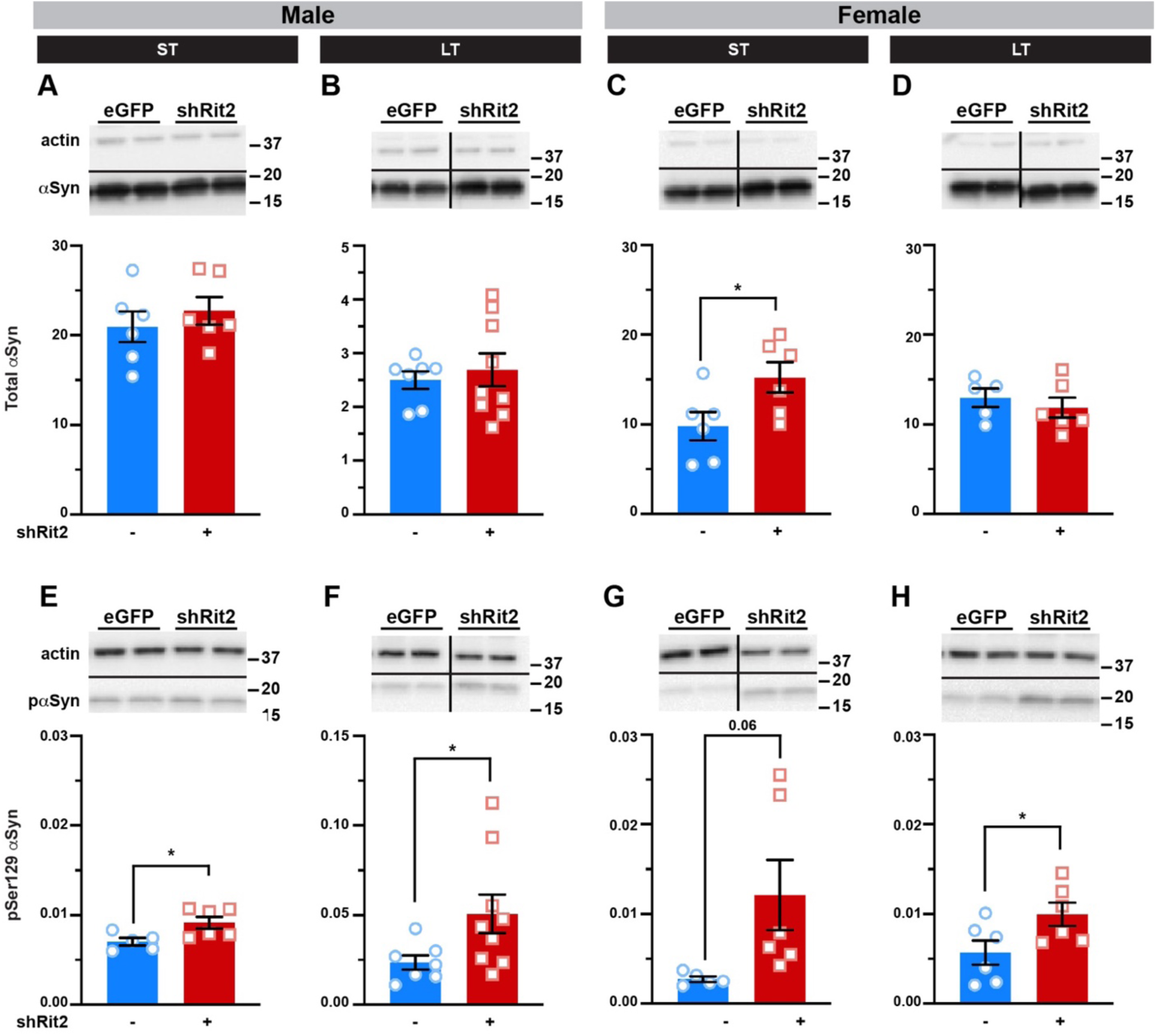
Prolonged DAergic Rit2 silencing increases PD-associated protein biomarkers in striatum. *Quantitative Immunoblotting.* Pitx3^IRES-tTA^ mouse VTA were bilaterally injected with either AAV9-TRE-eGFP or AAV9-TRE-shRit2. Striatal lysates were collected 4-5 weeks (ST) or 5-6mo (LT) post-injection and αSyn and pS129-αSyn levels were measured by quantitative immunoblot, normalized to actin, as described in *Methods*. *Tops:* Representative immunoblots showing two independent mouse lysates per virus. N values indicate the number of independent striata assessed. **(A-D)** α*Syn levels:* shRit2 had no effect on total αSyn levels in males at either ST (**A.** p=0.46, n=6) or LT (**B.** p=0.62, n=7-9) timepoints. ST shRit2 significantly increased αSyn in females (**C.** *p=0.04, n=6), but had no effect in LT females as compared to controls (**D.** p=0.50, n=6). (**E-H)** pS129-α*Syn levels*: pS129-αSyn was significantly increased by shRit2 in ST (**E. \***p=0.02, n=6) and LT (**F.** *p=0.04, n=7-9) males. pS129-αSyn trended towards a significant increase by shRit2 in ST females (**G.** p=0.06, n=6) and was significantly increased in LT females (**H.** *p=0.04, n=7-9). Two-tailed, unpaired, Student’s t test.

We additionally performed immunohistochemistry to ask whether LT Rit2 KD either 1) deleteriously affected DAergic terminals in the DS, or 2) impacted pSer129-Syn aggregation, which is a hallmark of PD. Both males and females exhibited a gross loss of TH+ terminals in DS, with little/no TH immunoreactivity detected (Fig. 5 G, H). Interestingly, in control mice pSer129 presented as both diffuse and punctate signals, whereas in shRit2 mice there was no apparent diffuse signal, with abundant puncta throughout the DS.

### Male motor learning is rescued with Parkinson’s therapeutics at early, but not late timepoints

The most widely used treatment strategy for PD is to increase DA availability by providing the DA precursor, L-DOPA. Moreover, recent studies suggest that increasing extracellular DA by inhibiting DAT with methylphenidate (Ritalin), may have therapeutic potential in PD^38,39^. We asked whether such pharmacological intervention could rescue the motor deficits observed on the rotarod due to Rit2 silencing. We first tested whether increasing extracellular DA levels by inhibiting DAT with methylphenidate would rescue motor learning. ST shRit2 males were assessed on the accelerating rotarod, injected with either saline or methylphenidate (MPH, 5mg/kg, I.P.), and were reassessed 15 min post-injection (see schematic, Figure 7A). MPH treatment significantly improved rotarod performance as compared to saline-injected mice (Figure 7B). MPH is equipotent at DAT and the norepinephrine transporter (NET)^40^. Therefore, to rule out any adrenergic contributions to the observed motor learning rescue, we tested whether rotarod performance was improved with desipramine (DMI), a NET-specific inhibitor. DMI treatment had no significant effect on shRit2 mouse performance (Figure 7B), suggesting that DAT inhibition was specifically responsible for rescued rotarod performance in ST shRit2 males. L-DOPA is the immediate chemical precursor to DA and is the prevailing PD treatment^41^. Therefore, we next asked whether L-DOPA treatment could rescue motor learning deficits in ST and LT shRit2 mice. Mice were scored on the accelerating rotarod, injected with either saline or L-DOPA (20mg/kg, I.P.), and rescored 1-hour post-injection. In ST shRit2 mice, L-DOPA significantly improved rotarod performance (Figure 7C). However, L-DOPA treatment had no effect on rotarod performance in LT shRit2 mice, (Figure 7D). Taken together, these data demonstrate that while pharmacological intervention can rescue motor deficits exhibited by ST shRit2 mice, the loss of DAergic tone and DANs caused by LT Rit2 KD drives deficits that are not rescuable by pharmacological means.

**Figure 7.**
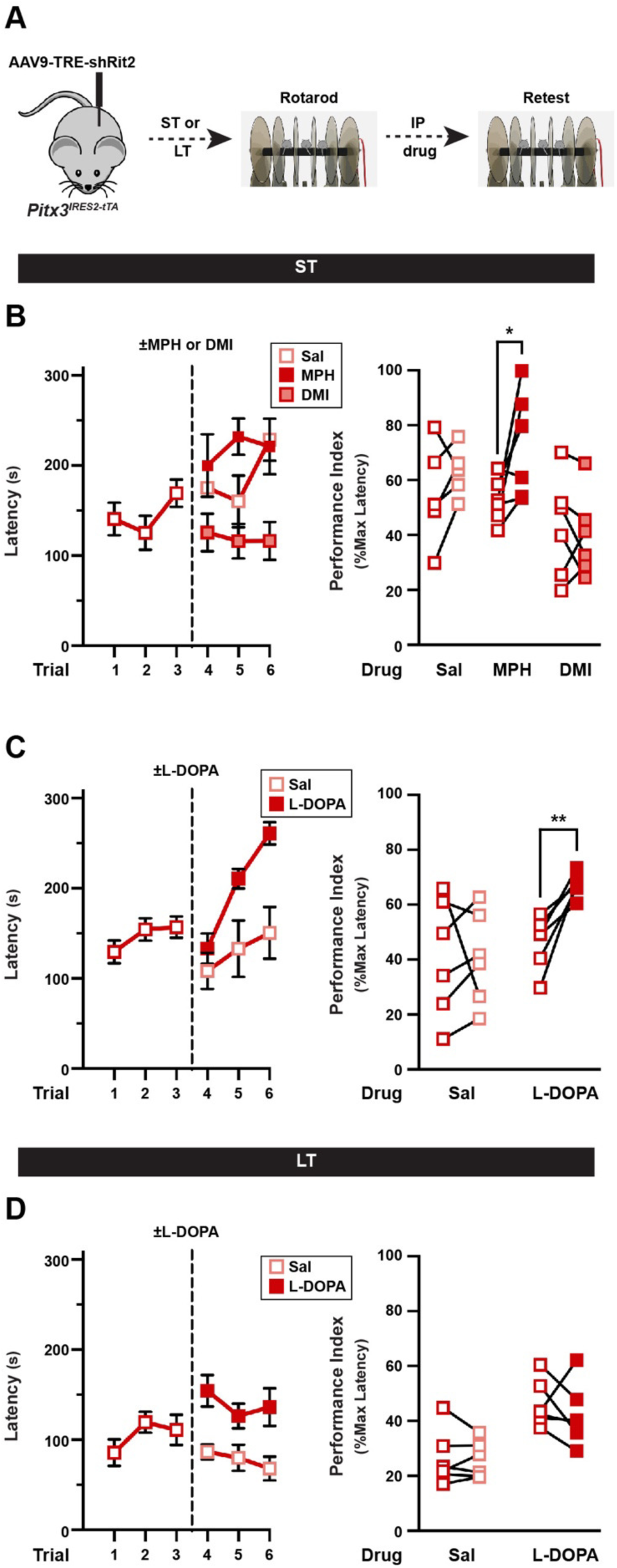
Male motor learning deficits are rescuable with Parkinson’s therapeutics at early, but not late, Rit2 silencing timepoints. *Accelerating rotarod rescue studies.* **(A)** *Experimental schematic.* Pitx3^IRES-tTA^ male mouse VTA were bilaterally injected with AAV9-TRE- shRit2 and were assessed on the accelerating rotarod at the indicated timepoints as described in *Methods.* Mice were initially tested for three trials, injected ±the indicated treatment drugs (I.P), and were retested for an additional three trials. Performance indices for each mouse were calculated pre- and post-test, and performance were assessed with a two-tailed, paired Student’s t test. N values indicate the number of independent mice. **(B)** *Methylphenidate (MPH) treatment:* ST shRit2 mice received either saline, 5mg/kg MPH, or 0.5mg/kg DMI, and were retested 15 min post-injection. *Left:* Raw rotarod results presented as latency to fall during the trial. *Right:* Paired pre- and post-test rotarod performance indices. shRit2 mice treated with MPH performed significantly better than pre-injection (*p=0.03), whereas performance was not enhanced by either saline (p=0.28) or DMI (p=0.60), n=5-6. **(C)** *L-DOPA treatment on ST shRit2 males:* ST shRit2 mice were treated ±20mg/kg L-DOPA and were retested 1 hour post-injection. *Left:* Raw rotarod results. *Right:* Paired pre- and post-test rotarod performance indices. shRit2 mice treated with L-DOPA (**p=0.005), but not saline (p=0.98) performed significantly better than pre-injection, n=6-7. **(D)** *L-DOPA treatment on LT shRit2 males:* LT shRit2 mice were treated ±20mg/kg L-DOPA and were retested 1 hour post-injection. *Left:* Raw rotarod results. *Right:* Paired pre- and post-test rotarod performance indices. Neither L-DOPA (p=0.48) nor saline (p=0.82) treatment significantly improved performance as compared to pre-injection performance, n=6.

## Discussion

Multiple GWAS studies report Rit2 as a PD risk allele^16,17,19–22,24,26^. Additionally, Rit2 mRNA is among the more highly downregulated genes in PD patient substantia nigra^3^, and defines a specific transcriptomic cluster in single-cell RNAseq studies from PD patients^32^. Despite these findings, it was unknown whether diminished Rit2 levels are causal or consequential for PD symptoms. Our results demonstrate that prolonged, conditional Rit2 silencing in DANs leads to PD-like phenotypes, both at the molecular and behavioral level. Specifically, DAergic Rit2 KD resulted in motor dysfunction (Figure 1), accompanied by decreases in DA release (Figure 2), striatal DA content (Figure 3), DAergic gene and protein expression (Figure 4), as well as decreased DAN numbers and striatal arborization (Figure 5). These were accompanied by increased pSer129-αSyn (Figure 6) and pSER935-LRRK2 (Supplemental Figure 6). Thus, our results demonstrate that Rit2 loss itself is causal for PD-like phenotypes.

In most instances, males were affected at an earlier timepoint than females, consistent with sex- specific differences in PD prevalence and onset in patients. Interestingly, female performance on the accelerating rotarod was completely resistant to Rit2 loss, even at the LT timepoint, despite significant deficits in gait, fix-speed rotarod, and challenge balance beam, and diminished dopaminergic tone. Motor learning on the accelerating rotarod is DA-dependent^42,43^, however the observed sexual dimorphism raises the possibility that there may be a strong DA- independent component to motor learning in females. Alternatively, it is possible that distinct, sex-specific SNc DAN subpopulations are vulnerable following Rit2 silencing. Future single cell transcriptomic studies will be necessary to test this possibility directly.

While motor learning deficits are specific to shRit2 male mice, shRit2 coordination deficits are sex-independent and are only apparent with prolonged Rit2 silencing (Figure 1). Coordination deficits were accompanied by modest alterations in shRit2 mouse gait (Supplemental Figure 2). Furthermore, LT male and female mice both exhibited increased four-limb grip strength (Supplemental Figure 2). While grip strength typically decreases with PD progression in patients^5,44^, increased mouse grip strength may also reflect rigidity or bradykinesia present in PD. Indeed, increased grip strength is observed in a rat 6-OHDA lesion PD model^45^.

Surprisingly, despite coordination deficits, conditional Rit2 silencing did not perturb horizontal locomotion, even with prolonged silencing (Supplemental Figure 1). PD is a late onset neurodegenerative disorder and motor symptoms are often not overtly apparent until >75% of SNc DANs have died^3,4^. Indeed, our stereological data suggest that at the LT timepoint (∼25 weeks post-injection), there is ∼32% loss of TH+ neurons in the SNc, and a gross loss of TH+ terminals in DS (Figure 5). Thus, it is possible that longer Rit2 silencing would lead to even further losses in the TH+ population, and more pronounced baseline motor deficits. Of note, the loss of TH+ striatal terminals is grossly disproportionate to the cell number loss in SNc (Figure 5). Recent studies indicate that DAergic axons likely deteriorate at a faster pace than their respective soma^46^, which may explain our observations

While Rit2 silencing diminished DAN numbers, the mechanism(s) downstream of Rit2 loss that led to decreased DAN viability are unknown. To date, the function of Rit2 in neurons remains poorly defined. Rit2 is required for EGF- and NGF-mediated neurite outgrowth in cell culture models^47^, and is required for NGF-mediated ERK phosphorylation and cell viability^13,14^. Our lab previously reported that Rit2 binds directly to the DAT^10^ and is required for both PKC- and mGluR5-stimulated DAT endocytosis in *ex vivo* striatal slices^8,9^. Indeed, ST DAergic mGluR5 silencing in male mice blocked DAT internalization, increased DAT plasma membrane presentation, and likewise drove motor learning dysfunction on the accelerating rotarod that was rescuable with a DAT-selective inhibitor^9^. Thus, ST shRit2 effects may be due, in part, to DAT dysregulation, while LT shRit2 effects are more likely due to diminished DAN viability. We also previously reported that ST DAergic Rit2 KD differentially modulates acute cocaine locomotor responses in males and females, wherein male shRit2 mice exhibit increased cocaine sensitivity, and females exhibit decreased cocaine sensitivity^34^. Thus, the requirement for Rit2 in DANs likely extends beyond motor behaviors.

Despite ST Rit2 KD significantly decreasing DAT protein levels (Figure 4J), there was no significant impact on DA clearance in parallel FSCV studies (Figure 2D). Interestingly, we instead observed that Rit2 silencing dampened DA release as compared to controls (Figure 2C). Decreased DA release was accompanied by decreases in both pSer40-TH (Supplemental Figure 3A) and DRD2 mRNA (Supplemental Figure 4A), raising the possibility that DA synthesis may be altered due to Rit2 silencing. However, DRD2-mediated regulation of DA clearance remained intact following Rit2 KD (Figure 2D), suggesting that DRD2 dysregulation is not likely responsible for the observed decrease in DA release. Rit2 may play some previously undefined role in DA synthesis and/or release that is independent of DRD2 regulation. Interestingly, DA release is adversely affected prior to DAN degeneration in several experimental PD models^46,48^, raising the possibility that the altered DA release we observed following ST Rit2 KD may, likewise, forbode the imminent DAN degeneration.

We also found that LT Rit2 KD significantly diminished expression of several hallmark DAergic genes, including TH, DAT, DRD2, and Pitx3, that are critical for DA signaling and the DAergic phenotype. In contrast, the DAN-specific transcription factor, Nurr1, was only decreased in LT shRit2 mice (Supplemental Figure 4). Nurr1 is associated with PD progression and DAergic Nurr1 ablation results in decreased, DAT and TH expression, reduced striatal DA content and locomotor deficits^49,50^. Surprisingly, the ubiquitously expressed genes, Rit1 and Vps35 were upregulated (Supplemental Figure 5). Rit1 is the closest homolog to Rit2 and may be upregulated or stabilized to compensate for Rit2 loss, however, whether they functionally overlap is unknown. Vps35 is a core retromer protein required for the endocytic delivery of DAT, and several neuronal receptors, to the plasma membrane^9,51–53^, and Vps35 mutations have been identified in PD patients. Thus, increased Vps35 expression may also be a compensatory mechanism to deliver additional DAT and/or receptors to the membrane in response to Rit2 loss. DAergic Rit2 KD also drove an increase in Ser129-αSyn, a PD biomarker (Figure 6). pSer129-αSyn accumulates in DAergic nuclei and negatively regulates Nurr1 expression^54^. Whether Rit2 directly regulates DAergic gene expression, or whether the observed changes are consequences of viability and cell death will need to be determined in future studies.

We used two pharmacological approaches to test whether boosting DA availability could rescue motor dysfunction at ST and LT timepoints: DAT inhibition with MPH, and L-DOPA treatment. Both MPH and L-DOPA treatments rescued male accelerating rotarod performance following ST Rit2 KD (Figure 7B,C). We previously reported that a DAT inhibitor (CE-158) can rescue motor learning dysfunction in response to DAergic mGluR5 KD, but has no effect on wildtype mice^9^. Thus, MPH-mediated rotarod rescue is likewise not likely to generally stimulate locomotion. While L-DOPA rescued ST motor learning deficits, it was unable to rescue motor performance following LT Rit2 KD (Figure 7D). PD patients typically lose responsiveness to L-DOPA during disease progression, and L-DOPA efficacy requires DA uptake and release mechanisms to be in place^55^. Given the striking loss of TH+ terminals in striatum (Figure 5), it is therefore not surprising that L-DOPA lost its efficacy at the LT timepoint.

Our study, for the first time demonstrates that DAergic Rit2 is required for DA neuron viability and DA-dependent motor behaviors. We identified a sex-dependent role for Rit2 in motor learning and demonstrated the manifestation of a PD-like phenotype, that included gene expression, DA content, and biomarker changes in DANs following prolonged Rit2 silencing. Future studies will illuminate the mechanisms that lead from Rit2 silencing to decreased DAN viability.

## Methods

### Materials

L-DOPA (3788) and L-741,626 (1003) were from Tocris. Methylphenidate and desipramine were from Sigma. All other reagents were from either Sigma-Aldrich or Fisher Scientific and were of the highest possible grade.

### Mice

*Pitx3^IRES-iTA/+^* mice were continuously backcrossed onto the C57Bl/6J background and were the generous gift of Dr. Huaibin Cai (National Institute on Aging). Mice were maintained on 12hr light/dark cycle (lights on at 0700) at constant temperature and humidity. Food and water were available ad libitum and mice were maintained on standard chow. All studies were conducted in accordance with University of Massachusetts Medical School IACUC Protocol 202100046 (formerly A-1506 H.E.M.).

### AAVs and Stereotaxic Surgeries

#### AAVs

pscAAV-TRE3g-eGFP and pscAAV-TRE3g-miR33-shRit2-eGFP AAV9 particles were produced as previously described^34^ by the University of Massachusetts Medical School Viral Vector Core.

#### Survival Surgeries

Mice aged 3-4 weeks were anesthetized with I.P. 100mg/kg ketamine (Vedco Inc.) and 10mg/kg xylazine (Akorn Inc). To increase viral spread, mice were administered 20% mannitol (NeogenVet) at least 15min prior to viral delivery^56^. Anesthetized mice were prepared and placed in a stereotaxic frame (Stoelting Inc.). 1µL of the indicated viruses were administered at a rate of 0.2µL/min bilaterally to the VTA using coordinates from bregma: anterior/posterior: - 3.08mm, medial/lateral: ±0.5mm, dorsal/ventral: -4.5mm. Syringes were maintained in position for a minimum of 5 minutes post-infusion prior to removal. Viral incubation was for either 4-5 weeks or 5-6months. Viral expression was confirmed by visualizing midbrain GFP reporters encoded in the viral constructs and/or by RT-qPCR.

### RNA Extraction and RT-qPCR

Bilateral 1.0mm^2^ tissue punches were obtained from 300µm coronal ventral midbrain slices of experimental mice. Punches were collected while visualizing GFP on an inverted fluorescence microscope and RNA was extracted immediately, or following tissue storage at -70°C, using RNAqueous®-Micro Kit RNA isolation (Thermo Fisher Scientific). Extracted RNA was reverse transcribed using RETROscript® reverse transcription kit (Thermo Fisher Scientific). Quantitative PCR was performed using the Applied Biosystems® 7500 Real-Time PCR System Machine and software or using the Bio-Rad C1000 Touch Thermal Cycler with CFX96 Real- Time system and software using Taqman® gene expression assays for mouse Rit2 (Mm0172749_mH), TH (Mm00447557_m1), DAT (Mm00438388_m1), DRD2 (Mm00438541_m1), Pitx3 (Mm01194166_g1), Nurr1 (Nr4a2, Mm00443060_m1), Rit1 (Mm00501400_m1), Vps35 (Mm00458167_m1). All Ct values were normalized to internal GAPDH (Mm99999915_g1) expression levels, to determine ΔCt values. For linear comparisons, data were analyzed by comparing 2^-ΔCt^ values.

### Mouse Behavior

#### Locomotion

Mouse activity was monitored individually in photobeam activity chambers (San Diego Instruments) as previously described ^34^. Horizontal, vertical, and fine movements were measured in 5-minute binds for 90 minutes total.

#### Accelerating and Fixed-Speed rotarod

Mice were habituated to the testing room in home cage for >30min with ambient lighting and the rotarod (UgoBasile 47600) running at 4 RPM.

#### Accelerating rotarod

Mice were placed on the rod moving at 4 RPM and rod speed was increased linearly from 4 to 40 RPM over 5 minutes. Trials were terminated and latency determined by either triggering the strike plate during a fall or if the mouse made >1 consecutive passive rotation. For drug administration studies, performance indices were calculated as the average latency time to fall (sec) across three trials (pre- or post-drug administration) normalized to the maximal trial time (300 sec). *Fixed-speed*: Mice were placed on the rod moving at the indicated speeds (20, 25, 30, 35, 40, 45 RPM) for 60 second trials. Latency to fall was measured or trial was stopped following >1 passive rotation. Two consecutive trials were performed per speed and latencies were averaged per animal.

#### Challenge Balance Beam

Mice were habituated to the testing room for >30min with overhead lights off and only a single light source placed approximately 1.5 feet over the beam origin illuminated. On day one, mice were trained over 5 trials to traverse a 1.0m, step-wise tapered (widths: 35mm, 25mm, 15mm, 5mm) elevated beam (#80306, Lafayette Neuroscience) at an incline of 15°. A dark box with home-cage bedding was placed at the far end of the beam. On day two, a challenge grid with 1cm x 1cm openings (custom 3D-printed, Thingiverse #4869650) was placed over the beam and mice traversed the beam in 3 independent trials. Traversal initiation and completion were determined by breaking an IR beam at each end of the beam. Traversals were video captured and scored for foot faults and traversal time, averaged over the first two completed trials. Both the experimenter and an independent scorer were double-blinded to the mouse ID.

#### Gait Analysis

Gait analysis protocol was adapted from Wertman, *et al*.^57^. Gait testing apparatus consisted of a 10cm x 36cm runway with 14cm high foamboard walls and a dark box at the opposing end. Fresh, legal-size paper was placed on the benchtop under the runway for each trial. Mouse forepaws and hind-paws were dipped in non-toxic orange and blue tempera paint, respectively, and mice were placed on the paper at the open end of the runway and allowed to traverse to the closed box at the opposing end. Three trials were performed per mouse and stride length, stride width and tow spread were measured for both fore- and hindlimbs. Number of completed trials was also quantified. Experimenters and data analysts were double-blinded to mouse IDs.

#### Grip Strength

Four-limb grip strength was measured using the Bioseb Grip Strength Test (BIO- GS3) equipped with mesh grip grid for mice. Mice were suspended by tail over the mesh and lowered onto it until all 4 paws grasped the mesh. The mouse was then pulled backwards gently on the horizontal plane until it released from the mesh. The maximal force applied was recorded for 3 consecutive trials and averaged.

### Tissue harvesting and immunoblotting

Striata were collected by preparing 300µm coronal sections on a Vibratome as previously described^8,9^. Sections were collected through the entire striatum, dorsal and ventral striata were sub-dissected, and slices encompassing each region were pooled for each independent mouse. Tissue was lysed in RIPA buffer (10mM Tris, pH 7.4; 150mM NaCl; 1.0mM EDTA; 0.1% SDS, 1% Triton X-100, 1% Na deoxycholate) supplemented with protease inhibitors (1.0mM phenylmethylsulfonyl fluoride and 1.0g/mL each leupeptin, aprotinin, and pepstatin) and Phosphatase inhibitor cocktail V (EMD Millipore). Mechanical tissue disruption was also performed by triturating sequentially through a 200µL pipette tip, 22-, and 26- gauge tech tips and solubilized by rotating (30min 4°C). Insoluble material was removed by centrifugation (15min, 18K x g, 4°C). Lysate protein concentrations were determined by BCA protein assay (Thermo Fisher Scientific). Protein samples were denatured in an equal volume of 2x Laemmli sample buffer and were either rotated (30min, RT) for membrane protein immunoblots or boiled (5min) for soluble protein immunoblots. Proteins were resolved by SDS-Page, transferred to nitrocellulose membranes, and the indicated proteins were detected and quantified by immunoblotting with the following antibodies: rat anti-DAT (MAB369, Millipore; 1:2000), rabbit anti-TH (AB152, Millipore, 1:10000), rabbit anti-pSer40 TH (AB5935, Millipore, 1:5000), rabbit anti-αSyn, rabbit anti-pSer129-αSyn, anti-LRRK2, anti-pSer935 LRRK2, mouse anti-actin (Santa Cruz, 1:5000). Secondary antibodies conjugated to horseradish peroxidase were all from Jackson ImmunoResearch and immunoreactive bands were visualized by chemiluminescence using SuperSignal West Dura (Thermo Scientific). Immunoblotting solutions were prepared in either PBS-T, or TBS-T (137mM NaCl, 2.7mM KCl, 19mM Tris base, ph7.4, 0.1% Tween20) when probing for phosphoproteins. Non-saturating immunoreactive bands were detected using either VersaDoc 5000MP or Chemidoc imaging stations (Bio-Rad) and were quantified using Quantity One software (Bio-Rad). Representative blots shown for a given condition were cropped from the same exposure of the same immunoblot and spliced together for presentation purposes only. Splice margins are indicated with a line. Brightness and contrast settings were identical for all immunoblot images presented.

### Fast-Scan Cyclic Voltammetry

Mice were sacrificed by cervical dislocation and rapid decapitation. Heads were immediately submerged in ice-cold NMDG cutting solution, pH 7.3-7.4 (20mM HEPES, 2.5mM KCl, 1.25mM NaH_2_PO_4_, 30mM NaHCO_3_, 25mM glucose, 0.5mM CaCl_2_·4H_2_O. 10mM MgSO_4_·7H_2_O, 92mM N-methyl-D-glucamine, 2mM thiourea, 5mM Na^+^-ascorbate, 3mM Na^+^-pyruvate). Brains were removed, glued to the stage of a VT1200S Vibroslicer (Leica) and submerged in ice-cold, oxygenated cutting solution. 300µm slices were prepared and were hemisected along the midline prior to recovering in ACSF (125mM NaCl, 2.5mM KCl, 1.24mM NaH_2_PO_4_, 26mM NaHCO_3_, 11mM glucose, 2.4mM CaCl_2_·4H_2_O,1.2mM MgCl_2_·6H_2_O, pH 7.4) at 31°C for a minimum of 1 hour prior to recording. Hemislices were moved to the recording chamber and were perfused with oxygenated ASCF supplemented with 500µM Na-Ascorbate. Glass pipettes containing a 7µm carbon-fiber microelectrode were prepared and preconditioned in ASCF by applying triangular voltage ramps (−0.4 to +1.2 and back to −0.4 V at 400 V/s), delivered at 60Hz for 1 hour. Recordings were performed at 10Hz. Electrodes were calibrated to a 1µM DA standard prior to recording. Electrodes were positioned in DS and DA transients were electrically evoked with a 250µA rectangular pulse every 2 min, using a concentric bipolar electrode placed ∼100µm from the carbon fiber electrode. Data were collected with a 3-electrode headstage, using an EPC10 amplifier (Heka) after low-pass filter at 10 kHz and digitized at 100 kHz, using Patchmaster software (Heka). A stable baseline was achieved after evoking six consecutive DA transients, after which experimental data were collected. Each biological replicate is the average of three evoked DA transients/slice, and a minimum of 3 independent mice were used to gather data from the indicated number of slices in each experiment. Data were analyzed in Igor Pro, using the Wavemetrics FSCV plugin (gift of Veronica Alvarez, NIAAA). Peak amplitudes were measured for each individual DA transient, and tau was calculated as 1/e according to the equation: y = y_0_ + A^((x-x)/tau))^.

### Mass Spectrometry

#### Sample Preparation

Brains were harvested, 1.0mm coronal sections were prepared and bilateral 1.0mm^2^ punches were each taken from dorsal and ventral striata. Each bilateral pair was solubilized in 10µL internal standard solution (200µM 13C_4_-GABA and 1µM 2H_3_-DA in water with 500µM ascorbic acid and 0.1% formic acid) and 50µl ice-cold acetonitrile with 1% formic acid. Samples were vortexed twice for 0.5min with a 1 min incubation on ice between vortexing and were sonicated in an ice-water bath until tissue was completely disrupted. Samples were centrifuged (10 min, 16,000 x g) and supernatants were collected for LC/MS/MS analysis. A standard (STD) solution containing 200 µM GABA, 1 µM dopamine, 500 µM ascorbic acid and 0.1 % formic acid was also prepared.

#### LC/MS/MS

10µl samples were injected in triplicate into a Thermo Scientific Ultimate 3000 HPLC system on a SeQuant ZIC-cHILIC column (2.1 x 100 mm, 3 µm) with a ZIC-cHILIC guard column (2.1 x 20 mm, 5 µm), coupled with a Thermo Scientific TSQ Quantiva triple quadrupole mass spectrometer. The mobile phase was water with 0.1% formic acid (A) and acetonitrile (B), and the elution program was as follows: 0 min 25% A, 0.5 min 25% A, 4.5 min 45% A, 5.0 min 70% A, 8.0 min 70% A, 8.1 min 25% A, 12.0 min 25% A at 0.2 mL/min. Ionization was operated in the positive mode with the voltage of 4.2 kV. The parameters were set as follow: sheath gas, 35 Arb, aux gas, 15 Arb, vaporizer temperature, 250 °C, ion transfer tube temperature, 325 °C. Multiple reaction monitoring (MRM) was performed using a cycle time of 0.3 s, CID gas pressure of 1.5 mTorr, Q1 resolution (FWHM) of 0.7 and Q3 resolution (FWHM) of 0.7. The MRM transitions 104.1>87 (GABA), 108.1>91 (^13^C_4_-GABA), 154.1>91 (dopamine) and 157.1>93 (^2^H_3_-dopamine) were selected for quantification. All data was integrated and processed in Xcalibur (Version 2.2, Thermo Scientific).

### Stereological analysis and Immunohistochemistry/confocal microscopy

Briefly, mice were perfused and fixed with freshly made 4% paraformaldehyde (PFA) in PBS. Brains were removed immediately and fixed again in 4% PFA followed by equilibration in 30% sucrose in PBS. Midbrains were removed for stereological analysis, and forebrains were used for immunohistochemistry/confocal microscopy.

### Stereological analysis

SNc total and TH+ neurons were quantified as previously described^58^. Fixed brains were imbedded in the OCT-compound media (Sakura) and frozen in liquid nitrogen. 40 µm cryosections were prepared through the midbrain a Leica CM3050s cryostat, and were stored in an antifreeze media containing 30% ethylene glycol, 25% glycerol, and 5% phosphate buffer. For stereology counting, 1 in every 5 sections was selected with a random start and a total of 6 brain slices on average were used for each mouse for IHC labeling for TH, including DAB enhancement, followed by Cresyl violet staining to reveal all neurons. Substantia nigra pars compacta were imaged using a Zeiss Axioplan 2 microscope equipped with a 20X objective, and Stereo Investigator was used to estimate the total number of neurons in the region of interest using the following parameters: frame sizes, 150 X 150 µm; grid sizes, 250 X 250 µm; top guard zone height, 2 µm; and optical dissector height, 8 µm. These parameters yielded a coefficient of error <10% throughout the analysis. Total cell numbers measured were weighted to section thickness for each mouse and were averaged across each cohort. Investigators performing stereological counting were blinded to mouse identity.

### Immunohistochemistry/confocal microscopy

25µm coronal sections through the striatum were prepared using a microtome and were co-stained with rabbit anti-pSer129-Syn (Cell Signaling #23706; 1:500) and chicken anti-TH (Millipore #AB9702, 1:500), followed by staining with secondary antibodies preabsorbed against mouse (Alexa568-goat anti- rabbit, Alexa647-donkey anti-chicken, Jackson ImmunoResearch). Z-stacks in the dorsal striatum were acquired with a Zeiss 700 LSM scanning confocal microscope using 555nm and 647nm lasers, and were pseudocolored to red and green, respectively. Acquisition settings within each channel (pinhole size, digital gain, and laser strength) were identical across all samples. Z-stacks were imported into ImageJ software where a representative plane was chosen, channels were separated (for individual red and green images), and images exported as tiff files. Files were subsequently imported into Adobe Photoshop, and levels were adjusted identically across all images.

### Statistics

Data analysis was performed with GraphPad Prism software. All data were assessed for normality and nonparametric tests were applied if data distribution was non-Gaussian. Outliers in each data set were identified using either Grubb’s or Rout’s outlier tests, with a or Q values set at 0.05 or 5%, respectively, and were removed from further analysis. Significant differences between two values were determined using either a one-tailed, two-tailed, or paired Student’s t test, as indicated. Differences amongst more than two conditions were determined using one- way or two-way ANOVA, as appropriate, and significant differences among individual values within the group were determined by post-hoc multiple comparison tests, as described for each experiment.

## Data Availability

All data generated or analyzed during this study are included in this published article and its supplementary information files.

## Study Approval

All mouse studies were conducted in accordance with UMASS Chan Medical IACUC protocol PROTO202100046 (H.E.M).

## Supporting information

Supplemental Figures Kearney et al

## Acknowledgements

These studies were supported by R01 DA015169 (H.E.M.), R01DA035224 (H.E.M.) and the Parkinson’s Foundation Research Center (Z.Y).

## Author information

P.J.K, H.E.M., Y.Z., Y.T., S.A.S., and Z.Y, designed the studies; P.J.K, Y.Z., E.K, Y.T., and T.C. acquired data; P.J.K, H.E.M., R.G.P., and R.R.F. analyzed data; P.J.K, Z.Y, Y.Z., Y.T., S.A.S., and H.E.M. wrote the manuscript.

## Competing Interests

All authors declare no financial or non-financial competing interests.

## Notes

<u>Conflict-of-interest statement</u> The authors have declared that no conflict of interest exists.

### Competing Interest Statement

The authors have declared no competing interest.

### Summary of Updates

New data and revised text

## References

1 Hartmann, A. Postmortem studies in Parkinson’s disease. Dialogues Clin Neurosci 6, 281–293 (2004).

2 Dickson, D. W. Parkinson’s disease and parkinsonism: neuropathology. Cold Spring Harb Perspect Med 2, doi:10.1101/cshperspect.a009258 (2012).

3 Pringsheim, T., Jette, N., Frolkis, A. & Steeves, T. D. The prevalence of Parkinson’s disease: a systematic review and meta-analysis. Mov Disord 29, 1583–1590, doi:10.1002/mds.25945 (2014).

4 Lees, A. J. Unresolved issues relating to the shaking palsy on the celebration of James Parkinson’s 250th birthday. Mov Disord 22 **Suppl 17**, S327–334, doi:10.1002/mds.21684 (2007).

5 Jankovic, J. Parkinson’s disease: clinical features and diagnosis. J Neurol Neurosurg Psychiatry 79, 368–376, doi:10.1136/jnnp.2007.131045 (2008).

6 Tambasco, N., Romoli, M. & Calabresi, P. Levodopa in Parkinson’s Disease: Current Status and Future Developments. Curr Neuropharmacol 16, 1239–1252, doi:10.2174/1570159X15666170510143821 (2018).

7 Zhou, Q., Li, J., Wang, H., Yin, Y. & Zhou, J. Identification of nigral dopaminergic neuron-enriched genes in adult rats. Neurobiol Aging 32, 313–326, doi:10.1016/j.neurobiolaging.2009.02.009 (2011).

8 Fagan, R. R. et al. Dopamine transporter trafficking and Rit2 GTPase: Mechanism of action and in vivo impact. J Biol Chem 295, 5229–5244, doi:10.1074/jbc.RA120.012628 (2020).

9 Kearney, P. J. et al. Presynaptic Gq-coupled receptors drive biphasic dopamine transporter trafficking that modulates dopamine clearance and motor function. J Biol Chem 299, 102900, doi:10.1016/j.jbc.2023.102900 (2023).

10 Navaroli, D. M. et al. The plasma membrane-associated GTPase Rin interacts with the dopamine transporter and is required for protein kinase C-regulated dopamine transporter trafficking. J Neurosci 31, 13758–13770, doi:10.1523/JNEUROSCI.2649-11.2011 (2011).

11 Hoshino, M. & Nakamura, S. The Ras-like small GTP-binding protein Rin is activated by growth factor stimulation. Biochem Biophys Res Commun 295, 651–656, doi:10.1016/s0006-291x(02)00731-3 (2002).

12 Spencer, M. L., Shao, H., Tucker, H. M. & Andres, D. A. Nerve growth factor-dependent activation of the small GTPase Rin. J Biol Chem 277, 17605–17615, doi:10.1074/jbc.M111400200 (2002).

13 Hoshino, M. & Nakamura, S. Small GTPase Rin induces neurite outgrowth through Rac/Cdc42 and calmodulin in PC12 cells. J Cell Biol 163, 1067–1076, doi:10.1083/jcb.200308070 (2003).

14 Shi, G. X., Han, J. & Andres, D. A. Rin GTPase couples nerve growth factor signaling to p38 and b-Raf/ERK pathways to promote neuronal differentiation. J Biol Chem 280, 37599–37609, doi:10.1074/jbc.M507364200 (2005).

15 Latourelle, J. C., Dumitriu, A., Hadzi, T. C., Beach, T. G. & Myers, R. H. Evaluation of Parkinson disease risk variants as expression-QTLs. PLoS One 7, e46199, doi:10.1371/journal.pone.0046199 (2012).

16 Pankratz, N. et al. Meta-analysis of Parkinson’s disease: identification of a novel locus, RIT2. Ann Neurol 71, 370–384, doi:10.1002/ana.22687 (2012).

17 Emamalizadeh, B. et al. RIT2, a susceptibility gene for Parkinson’s disease in Iranian population. Neurobiol Aging 35, e27–e28, doi:10.1016/j.neurobiolaging.2014.07.013 (2014).

18 Nalls, M. A. et al. Large-scale meta-analysis of genome-wide association data identifies six new risk loci for Parkinson’s disease. Nat Genet 46, 989–993, doi:10.1038/ng.3043 (2014).

19 Lu, Y. et al. Genetic association of RIT2 rs12456492 polymorphism and Parkinson’s disease susceptibility in Asian populations: a meta-analysis. Sci Rep 5, 13805, doi:10.1038/srep13805 (2015).

20 Liu, Z. H. et al. Assessment of RIT2 rs12456492 association with Parkinson’s disease in Mainland China. Neurobiol Aging 36, 1600 e1609–1611, doi:10.1016/j.neurobiolaging.2014.12.012 (2015).

21 Wang, J. Y. et al. The RIT2 and STX1B polymorphisms are associated with Parkinson’s disease. Parkinsonism Relat Disord 21, 300–302, doi:10.1016/j.parkreldis.2014.12.006 (2015).

22 Zhang, X., Niu, M., Li, H. & Xie, A. RIT2 rs12456492 polymorphism and the risk of Parkinson’s disease: A meta-analysis. Neurosci Lett 602, 167–171, doi:10.1016/j.neulet.2015.07.004 (2015).

23 Chan, G. et al. Trans-pQTL study identifies immune crosstalk between Parkinson and Alzheimer loci. Neurol Genet 2, e90, doi:10.1212/NXG.0000000000000090 (2016).

24 Emamalizadeh, B. et al. RIT2 Polymorphisms: Is There a Differential Association? Mol Neurobiol 54, 2234–2240, doi:10.1007/s12035-016-9815-4 (2017).

25 Chang, D. et al. A meta-analysis of genome-wide association studies identifies 17 new Parkinson’s disease risk loci. Nat Genet 49, 1511–1516, doi:10.1038/ng.3955 (2017).

26 Daneshmandpour, Y., Darvish, H. & Emamalizadeh, B. RIT2: responsible and susceptible gene for neurological and psychiatric disorders. Mol Genet Genomics 293, 785–792, doi:10.1007/s00438-018-1451-4 (2018).

27 Glessner, J. T. et al. Strong synaptic transmission impact by copy number variations in schizophrenia. Proc Natl Acad Sci U S A 107, 10584–10589, doi:10.1073/pnas.1000274107 (2010).

28 Liu, H., Talalay, P. & Fahey, J. W. Biomarker-Guided Strategy for Treatment of Autism Spectrum Disorder (ASD). CNS Neurol Disord Drug Targets 15, 602–613, doi:10.2174/1871527315666160413120414 (2016).

29 Hamedani, S. Y. et al. Ras-like without CAAX 2 (RIT2): a susceptibility gene for autism spectrum disorder. Metab Brain Dis 32, 751–755, doi:10.1007/s11011-017-9969-4 (2017).

30 Bouquillon, S. et al. A 5.3Mb deletion in chromosome 18q12.3 as the smallest region of overlap in two patients with expressive speech delay. Eur J Med Genet 54, 194–197, doi:10.1016/j.ejmg.2010.11.009 (2011).

31 Bossers, K. et al. Analysis of gene expression in Parkinson’s disease: possible involvement of neurotrophic support and axon guidance in dopaminergic cell death. Brain Pathol 19, 91–107, doi:10.1111/j.1750-3639.2008.00171.x (2009).

32 Wang, Q. et al. Single-cell transcriptomic atlas of the human substantia nigra in Parkinson’s disease. bioRxiv, 2022.2003.2025.485846, doi:10.1101/2022.03.25.485846 (2022).

33 Obergasteiger, J. et al. The small GTPase Rit2 modulates LRRK2 kinase activity, is required for lysosomal function and protects against alpha-synuclein neuropathology. NPJ Parkinsons Dis 9, 44, doi:10.1038/s41531-023-00484-2 (2023).

34 Sweeney, C. G. et al. Conditional, inducible gene silencing in dopamine neurons reveals a sex-specific role for Rit2 GTPase in acute cocaine response and striatal function. Neuropsychopharmacology 45, 384–393, doi:10.1038/s41386-019-0457-x (2020).

35 Ford, C. P. The role of D2-autoreceptors in regulating dopamine neuron activity and transmission. Neuroscience 282, 13–22, doi:10.1016/j.neuroscience.2014.01.025 (2014).

36 Sato, H., Kato, T. & Arawaka, S. The role of Ser129 phosphorylation of alpha-synuclein in neurodegeneration of Parkinson’s disease: a review of in vivo models. Rev Neurosci 24, 115–123, doi:10.1515/revneuro-2012-0071 (2013).

37 Nichols, R. J. LRRK2 Phosphorylation. Adv Neurobiol 14, 51–70, doi:10.1007/978-3-319-49969-7_3 (2017).

38 Casiraghi, A. et al. Methylphenidate Analogues as a New Class of Potential Disease- Modifying Agents for Parkinson’s Disease: Evidence from Cell Models and Alpha- Synuclein Transgenic Mice. Pharmaceutics 14, doi:10.3390/pharmaceutics14081595 (2022).

39 Lou, J. S. Fatigue in Parkinson’s disease and potential interventions. NeuroRehabilitation 37, 25–34, doi:10.3233/NRE-151238 (2015).

40 Eshleman, A. J. et al. Characteristics of drug interactions with recombinant biogenic amine transporters expressed in the same cell type. J Pharmacol Exp Ther 289, 877–885 (1999).

41 Fahn, S. The history of dopamine and levodopa in the treatment of Parkinson’s disease. Mov Disord 23 **Suppl 3**, S497–508, doi:10.1002/mds.22028 (2008).

42 Augustin, S. M., Loewinger, G. C., O’Neal, T. J., Kravitz, A. V. & Lovinger, D. M. Dopamine D2 receptor signaling on iMSNs is required for initiation and vigor of learned actions. Neuropsychopharmacology 45, 2087–2097, doi:10.1038/s41386-020-00799-1 (2020).

43 Lemos, J. C. et al. Enhanced GABA Transmission Drives Bradykinesia Following Loss of Dopamine D2 Receptor Signaling. Neuron 90, 824–838, doi:10.1016/j.neuron.2016.04.040 (2016).

44 Roberts, H. C. et al. The Association of Grip Strength With Severity and Duration of Parkinson’s: A Cross-Sectional Study. Neurorehabil Neural Repair 29, 889–896, doi:10.1177/1545968315570324 (2015).

45 Jeyasingham, R. A., Baird, A. L., Meldrum, A. & Dunnett, S. B. Differential effects of unilateral striatal and nigrostriatal lesions on grip strength, skilled paw reaching and drug- induced rotation in the rat. Brain Res Bull 55, 541–548, doi:10.1016/s0361-9230(01)00557-3 (2001).

46 Cheng, H. C., Ulane, C. M. & Burke, R. E. Clinical progression in Parkinson disease and the neurobiology of axons. Ann Neurol 67, 715–725, doi:10.1002/ana.21995 (2010).

47 Shi, G. X. & Andres, D. A. Rit contributes to nerve growth factor-induced neuronal differentiation via activation of B-Raf-extracellular signal-regulated kinase and p38 mitogen-activated protein kinase cascades. Mol Cell Biol 25, 830–846, doi:10.1128/MCB.25.2.830-846.2005 (2005).

48 Cramb, K. M. L., Beccano-Kelly, D., Cragg, S. J. & Wade-Martins, R. Impaired dopamine release in Parkinson’s disease. Brain 146, 3117–3132, doi:10.1093/brain/awad064 (2023).

49 Decressac, M., Volakakis, N., Bjorklund, A. & Perlmann, T. NURR1 in Parkinson disease--from pathogenesis to therapeutic potential. Nat Rev Neurol 9, 629–636, doi:10.1038/nrneurol.2013.209 (2013).

50 Kadkhodaei, B. et al. Transcription factor Nurr1 maintains fiber integrity and nuclear- encoded mitochondrial gene expression in dopamine neurons. Proc Natl Acad Sci U S A 110, 2360–2365, doi:10.1073/pnas.1221077110 (2013).

51 Wu, S. et al. The Dopamine Transporter Recycles via a Retromer-Dependent Postendocytic Mechanism: Tracking Studies Using a Novel Fluorophore-Coupling Approach. J Neurosci 37, 9438–9452, doi:10.1523/JNEUROSCI.3885-16.2017 (2017).

52 Choy, R. W. et al. Retromer mediates a discrete route of local membrane delivery to dendrites. Neuron 82, 55–62, doi:10.1016/j.neuron.2014.02.018 (2014).

53 Temkin, P. et al. The Retromer Supports AMPA Receptor Trafficking During LTP. Neuron 94, 74–82 e75, doi:10.1016/j.neuron.2017.03.020 (2017).

54 Jia, C. et al. alpha-Synuclein Negatively Regulates Nurr1 Expression Through NF- kappaB-Related Mechanism. Front Mol Neurosci 13, 64, doi:10.3389/fnmol.2020.00064 (2020).

55 Nonnekes, J. et al. Unmasking levodopa resistance in Parkinson’s disease. Mov Disord 31, 1602–1609, doi:10.1002/mds.26712 (2016).

56 Burger, C., Nguyen, F. N., Deng, J. & Mandel, R. J. Systemic mannitol-induced hyperosmolality amplifies rAAV2-mediated striatal transduction to a greater extent than local co-infusion. Mol Ther 11, 327–331, doi:10.1016/j.ymthe.2004.08.031 (2005).

57 Wertman, V., Gromova, A., La Spada, A. R. & Cortes, C. J. Low-Cost Gait Analysis for Behavioral Phenotyping of Mouse Models of Neuromuscular Disease. J Vis Exp, doi:10.3791/59878 (2019).

58 Wang, Q. et al. The landscape of multiscale transcriptomic networks and key regulators in Parkinson’s disease. Nature Communications 10, 5234, doi:10.1038/s41467-019-13144-y (2019).

