## Supplemental Figures Kearney et al for "Rit2 silencing in dopamine neurons drives a Parkinsonian phenotype"

### Supplementary Figures

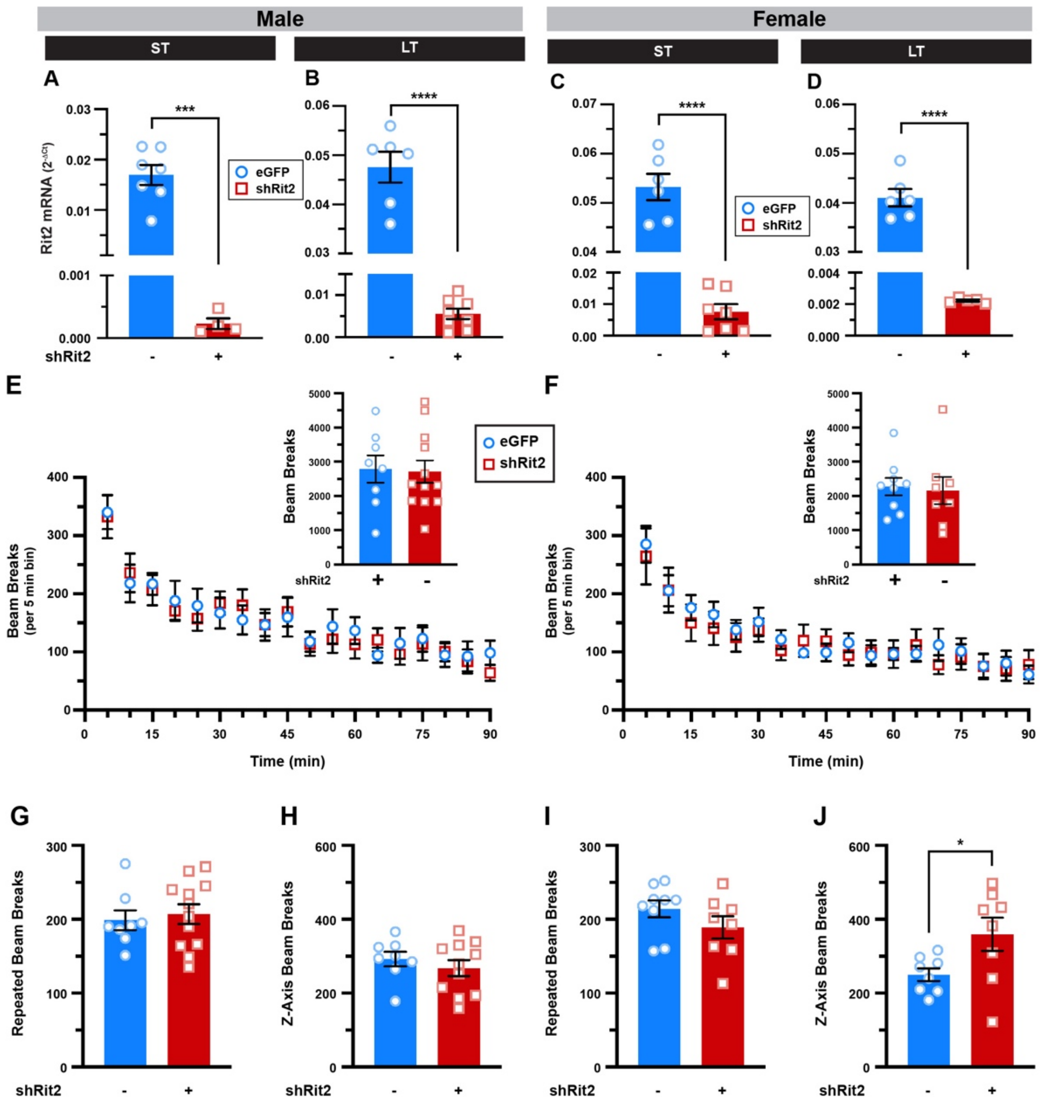


**Supplementary Figure 1. Effect of long-term conditional Rit2 silencing on mouse baseline locomotion. (A-D)** *Midbrain RT-qPCR. $Pitx3^{IRES\text{-}tTA}$* mouse VTA were bilaterally injected with either AAV9-TRE-eGFP or AAV9-TRE-shRit2 and midbrain punches were assessed for Rit2 expression by RT-qPCR at the indicated timepoints. Rit2 mRNA was significantly suppressed in both male and female mice at each timepoint. ST males (**A**, ***p=0.0002, n=7 (eGFP) n=4 (shRit2)), LT males (**B**, ****p< 0.0001, n=6 (GFP), n=8 (shRit2)), ST females (**C**, ****p< 0.0001. n=6 (eGFP), n=7 (shRit2)), LT females (**D**, ****p< 0.0001. n=6 (eGFP), n=5 (shRit2)). Two-tailed, unpaired Student's *t* test either with (**A,D)** or without (**B,C**) Welch's correction. *Mouse locomotor studies. $Pitx3^{IRES\text{-}tTA}$* mouse VTA were bilaterally injected with either AAV9-TRE-eGFP or AAV9-TRE-shRit2 and baseline locomotor activity was monitored in photobeam activity chambers after 5-6mo viral incubation as described in *Methods.* **(E,F)** *Total horizontal locomotion over time*. Conditional Rit2 silencing had no effect on male (**E**) or female (**F**) horizontal locomotion over 90min. *Insets: Averaged data.* Average beam breaks per session ±S.E.M. Conditional *Rit2* silencing had no significant effect on male (**E.** p= 0.89) or female (**F.** p= 0.81) horizontal locomotion. **(G-J)** *Fine and vertical movement*. Conditional Rit2 silencing had no effect on male (**G.** p=0.68) or female (**I.** p= 0.20) fine movement, or on male vertical movement (**H.** p= 0.43), whereas female vertical movement increased (**J.** *p=0.05). Two-tailed, unpaired, Student's or Welch's t test. males: n=8 (eGFP) and n=12 (shRit2), females: n=9 (eGFP) and n=8 (shRit2).

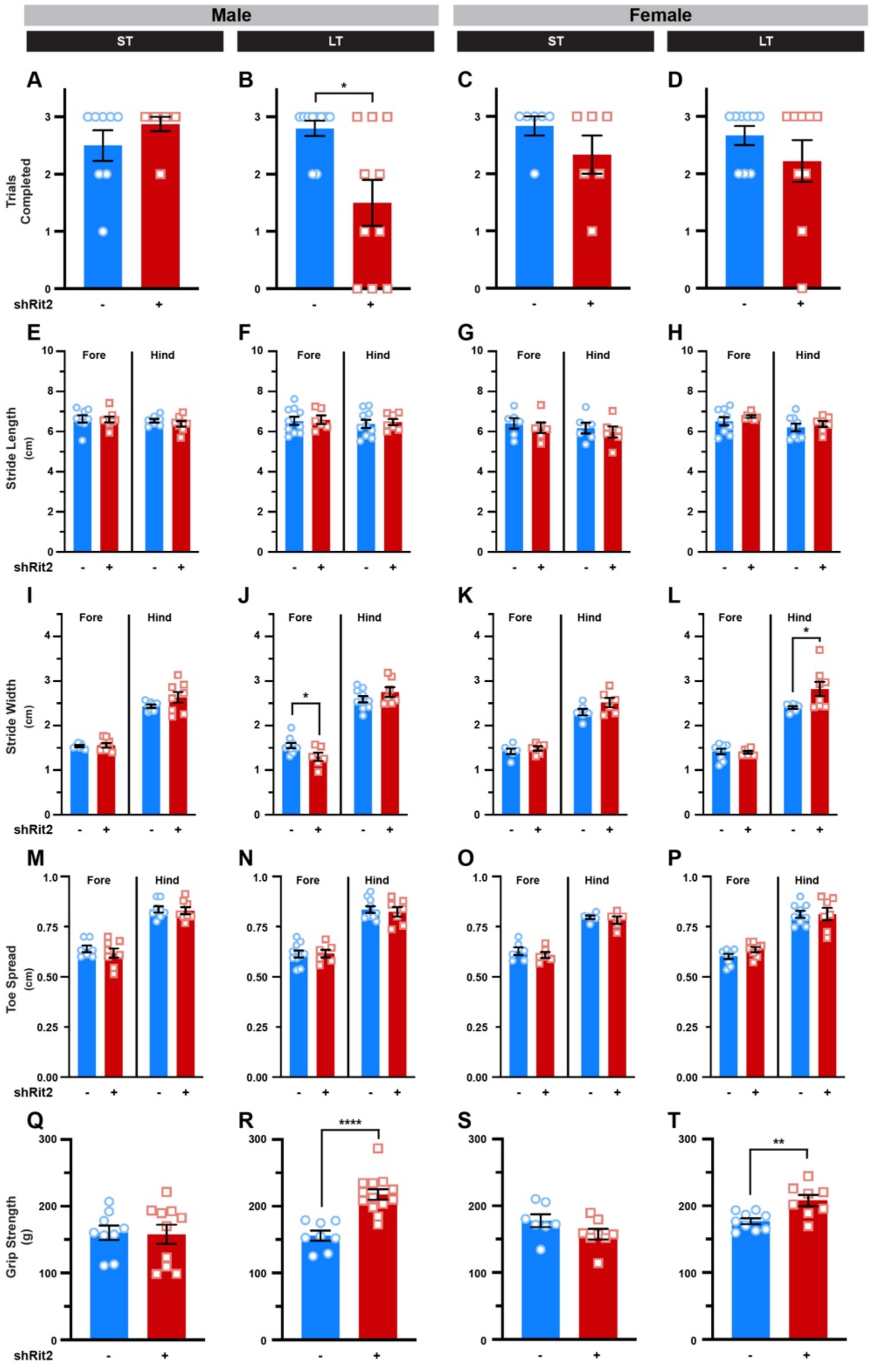


**Supplementary Figure 2. Effect of conditional *Rit2* silencing on mouse gait and grip strength.** *Pitx3*$^{IRES\text{-}tTA}$ mouse VTA were bilaterally injected with either AAV9-TRE-eGFP or AAV9-TRE-shRit2 and mouse gait and grip strength were assessed as described in *Methods.* Data were analyzed by an unpaired, two-tailed Student's t test. **(A-D)** *Gait analysis, trials completed:* There was no significant effect on ST (**A.** p= 0., n=) but significantly decreased trials completed for LT shRit2 males (**B.** *p=0.01 with Welch's correction, n=10**)** as compared to controls. Neither ST (**C.** p= 0., n=) nor LT shRit2 females (**D.** p= 0.29 with Welch's correction, n=9) were affected as compared to controls. **(E-H)** *Stride Length.* Forelimb and hindlimb stride length were not significantly affected by shRit2 in either ST males (**E.** fore: p= 0.91, hind: p=0.37, n=), LT males (**F.** fore: p= 0.85, hind: p=0.74, n=7-10), ST females (**G.** fore: p= 0.56, hind: p= 0.63, n=), or LT females (**H.** fore: p= 0.30, with Welch's correction, hind: p=0.50, n=7-9). **(I-L)** *Stride Width.* Forelimb and hindlimb stride width were not significantly affected by shRit2 in ST males (**I.** fore: p= 0.75, hind: p=0.16, n=), while only forelimb width was affected in LT males (**J.** fore: *p= 0.03, hind: p=0.21, n=7-10). In shRit2 females, there was no effect on ST females (**K.** fore: p= 0.45, hind: p= 0.12, n=), but hindlimbs were affected in LT females (**L.** fore: p= 0.81, hind: *p=0.04 with Welch's correction, n=7-9). **M-P.** *Toe Spread.* Forelimb and hindlimb toe spread were not significantly affected by shRit2 in either ST males (**M.** fore: p= 0.48, hind: p=0.80, n=), LT males (**N.** fore: p= 0.93, hind: p=0.69, n=6-10), ST females, (**O.** fore: p= 0.46, hind: p= 0.86, n=), or LT female hindlimb, and there was a trend for increased toe spread in LT female forelimb (**P.** fore: p= 0.08, hind: p=0.98, n=7-9). **(Q-T)** *Grip Strength.* Male shRit2 mouse grip strength was unaffected at the ST timepoint (**Q.** p=0.90, n=9-10), but significantly increased at the LT timepoint (**R.** ****p<0.0001, n=8-13) as compared to controls. Female shRit2 mouse grip strength was unaffected at the ST timepoint (**S.** p=0.12, n=7-8), but significantly increased at the LT timepoint (**T.** **p=0.004, n=8-9) as compared to controls.

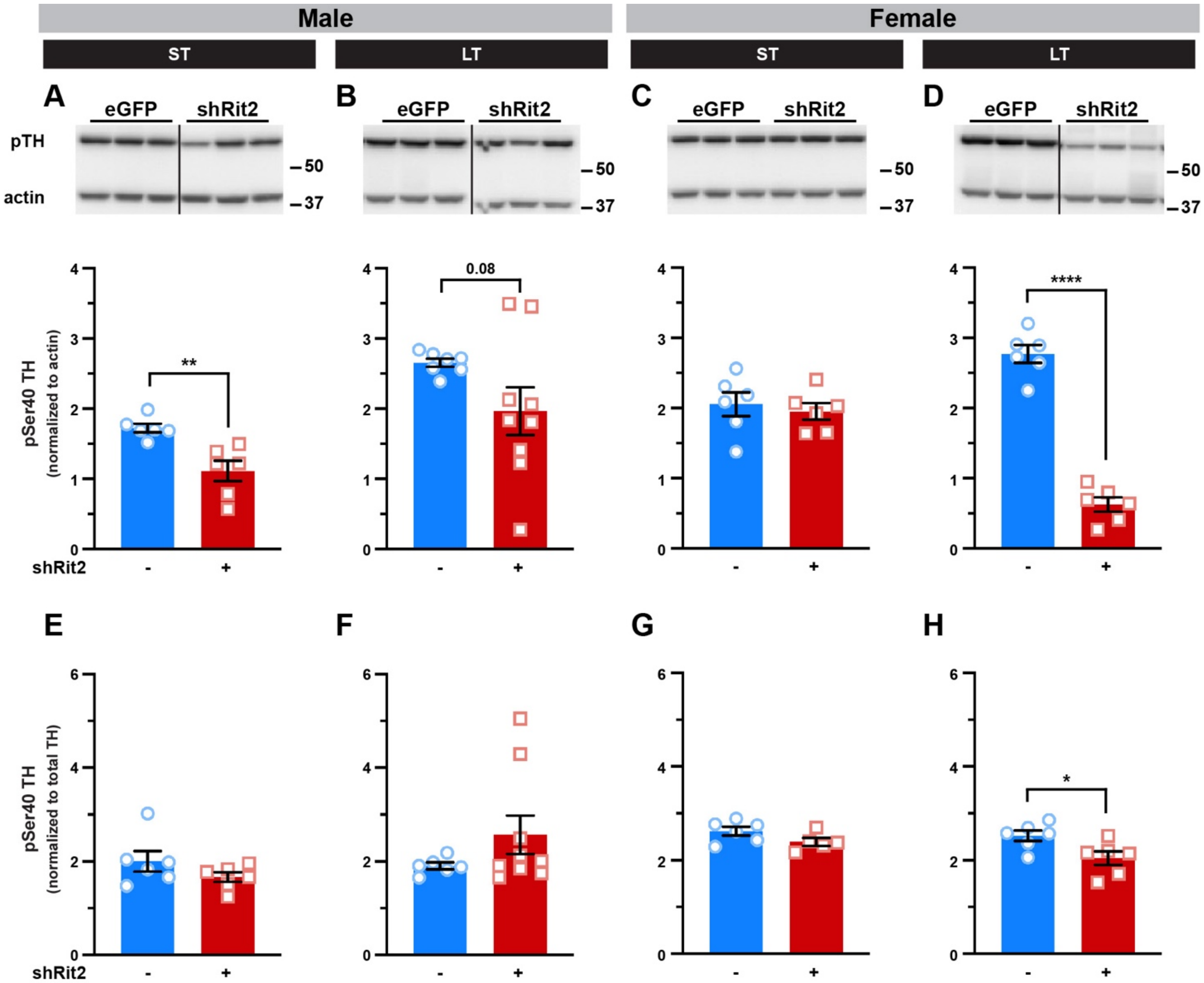


**Supplemental Figure 3. Effect of DAergic Rit2 silencing on striatal TH activation.** Male and female *Pitx3*$^{IRES\text{-}tTA}$ mouse VTA were bilaterally injected with either AAV9-TRE-eGFP or AAV9-TRE-shRit2 and striatum were dissected from male and female control and shRit2 mice at the indicated timepoints, lysed, and pSer40TH levels were assessed by quantitative immunoblot as described in *Methods*. Data were analyzed by unpaired, two-tailed Student's t test. N values indicate independent animals **(A-D)** *pSer40TH normalized to actin. Top:* Representative striatal immunoblots for each protein, showing 3 independent mouse lysates each for control (eGFP) and shRit2 mice. Molecular weight markers are indicated in kDa. pSer40-TH levels were decreased in ST shRit2 males (**A.** **p=0.003, n=6) and trended towards a decrease in LT shRit2 males (**B.** p=0.08 with Welch's correction, n=7-9) as compared to controls. pSer40-TH was unchanged in ST shRit2 females (**C.** p=0.63, n=6), but significantly decreased in LT shRit2 mice (**D.** ****p<0.0001, n=6). (**E-H**) *pSer40TH normalized to total TH.* pSer40TH levels from (A-D) were normalized to total TH from the same mice, presented in Figure 5. Fractional pSer40TH levels in ST (**E.** p=0.20, n=6) and LT (**F.** p=0.14 with Welch's correction, n=6-9) shRit2 males were not significantly affected as compared to controls. Fractional pSer40TH levels in ST shRit2 females were unaffected (**G.** p=0.11, n=5-6), but were significantly decreased in LT shRit2 females (**H.** *p<0.03, n=6) shRit2 males were not significantly affected as compared to controls.

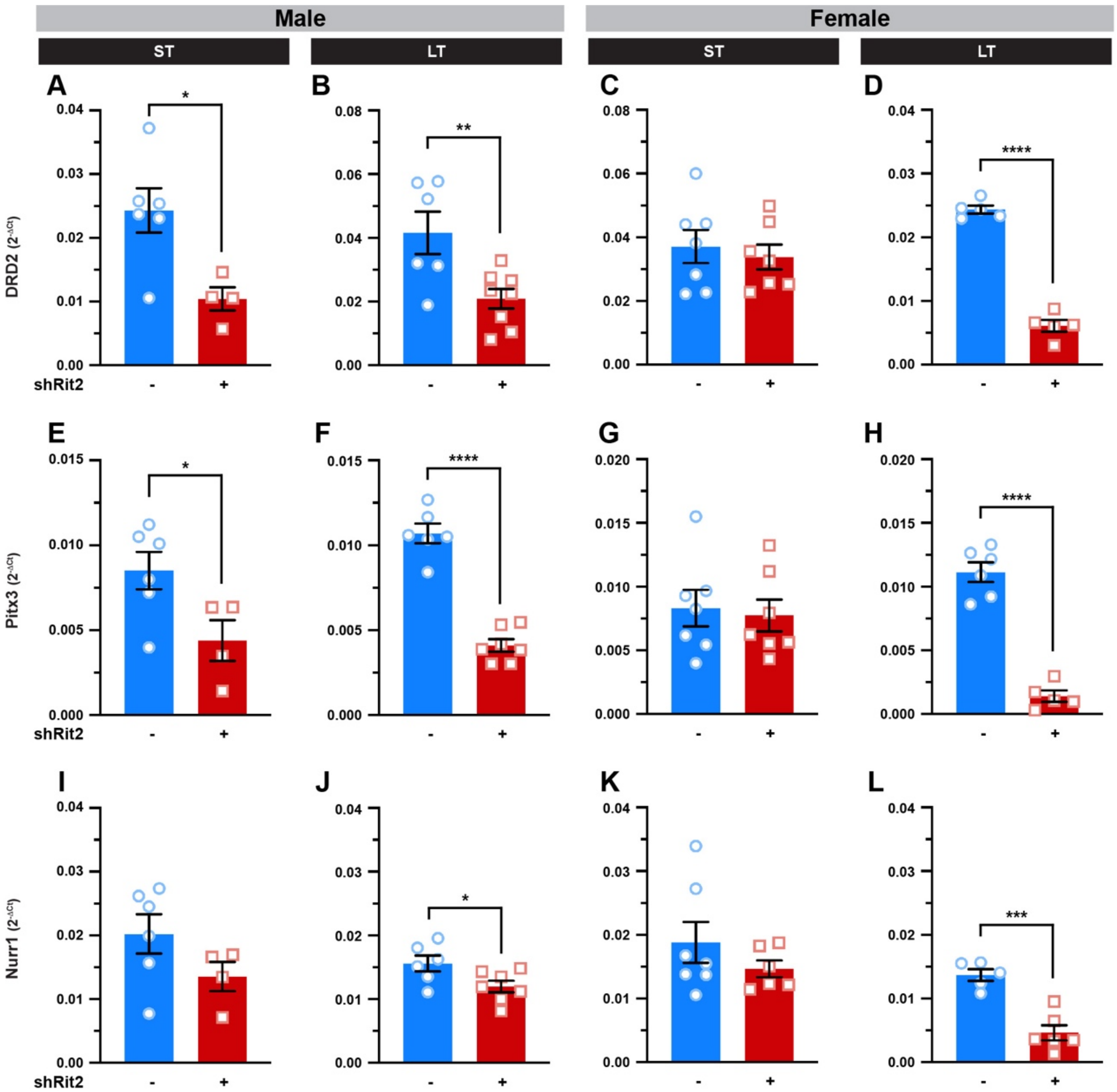


**Supplemental Figure 4. DAergic Rit2 silencing leads to a progressive decrease in DAergic gene expression.** *Ventral midbrain RT-qPCR studies.* Male and female *Pitx3*$^{IRES\text{-}tTA}$ mouse VTA were bilaterally injected with either AAV9-TRE-eGFP or AAV9-TRE-shRit2, midbrain tissue punches were harvested at ST and LT timepoints, and midbrain RNA levels were measured by RT-qPCR for the indicated mRNAs as described in *Methods*. Data were analyzed by two-tailed, unpaired, Student's t test. **(A-D)** *DRD2*: DRD2 gene expression was significantly decreased in ST (**A.** *p=0.02, n=4-6) and LT (**B.** **p=0.01, n=6-8) shRit2 male mice. In females, shRit2 had no effect on DRD2 expression at the ST timepoint (**C.** p=0.62, n=7), but significantly decreased DRD2 at the LT timepoint (**D.** ****p<0.0001, n=5). **(E-H)** *Pitx3*: Pitx3 gene expression was significantly decreased in both ST (**E.** *p=0.04, n=4-6) and LT (**F.** ****p<0.0001, n=6-7) shRit2 male mice. In females, shRit2 had no effect on DRD2 expression at the ST timepoint (**G.** p=0.76, n=7), but significantly decreased Pitx3 at the LT timepoint (**H.** ****p<0.0001, n=5-6). **(I-L)** *Nurr1*: Nurr1 gene expression was unchanged in ST shRit2 males (**I.** p=0.15, n=4-6) but was significantly decreased in LT shRit2 males (**J.** *p=0.04). In females, shRit2 had no effect on Nurr1 expression at the ST timepoint (**K.** p=0.29, n=6-7) but significantly decreased Pitx3 at the LT timepoint (**L.** ****p<0.0001, n=5).

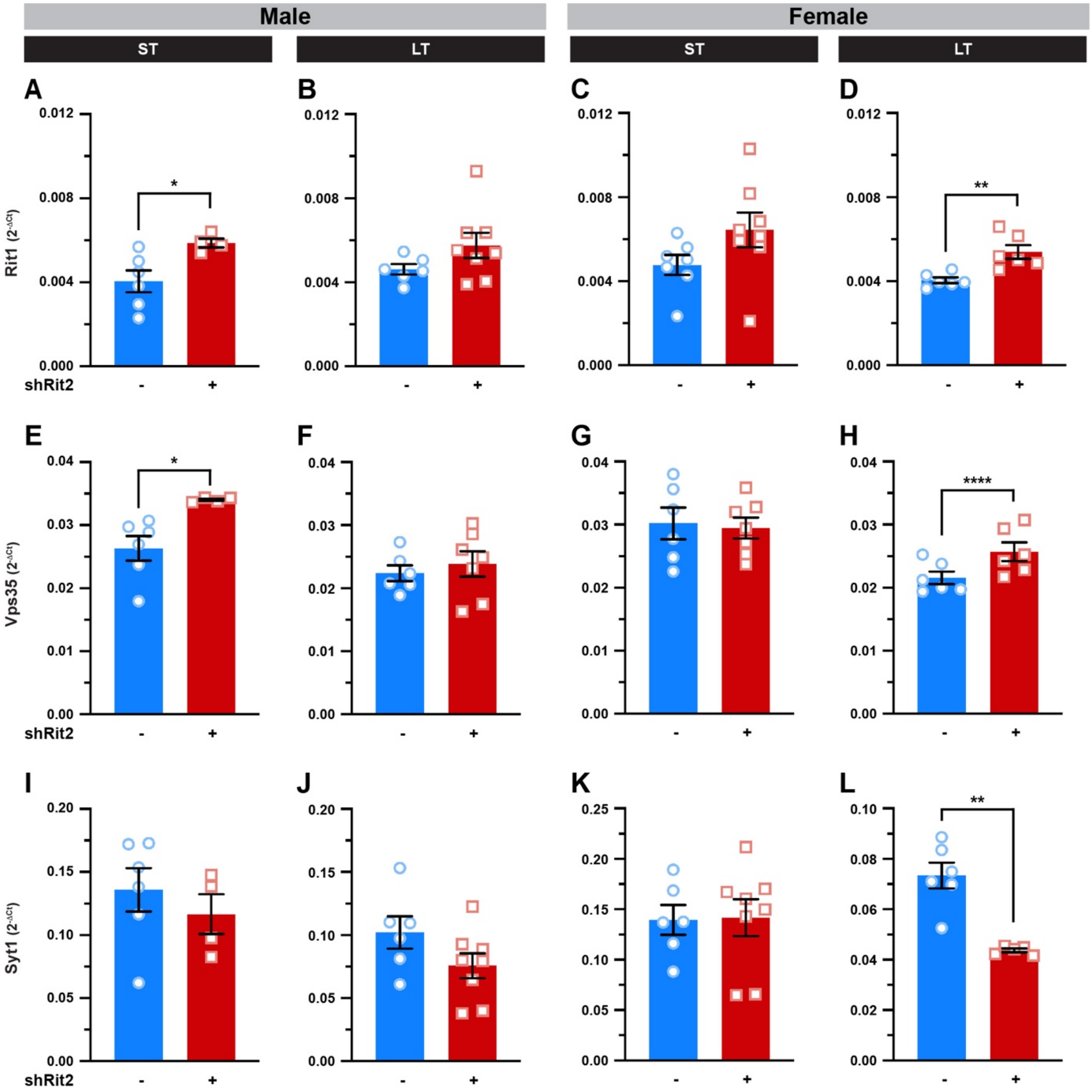


**Supplemental Figure 5. Effect of DAergic Rit2 KD on ubiquitous and pan neuronal ventral midbrain gene expression.** *Ventral midbrain RT-qPCR studies.* Male and female *Pitx3*$^{IRES\text{-}tTA}$ mouse VTA were bilaterally injected with either AAV9-TRE-eGFP or AAV9-TRE-shRit2, midbrain tissue punches were harvested at ST and LT timepoints, and midbrain RNA levels were measured by RT-qPCR for the indicated mRNAs as described in *Methods*. Data were analyzed by two-tailed, unpaired, Student's t test. **(A-D)** *Rit1*: In ST shRit2 males, Rit1 was transiently increased (**A.** *p=0.03, n=4-6), but was unchanged in LT shRit2 male mice (**B.** p=0.11 with Welch's correction, n=6-8). In females, Rit1 was unchanged at the ST timepoint (**C.** p=0.11, n=7-8), but was significantly increased at the LT timepoint (**D.** **p=0.006, n=5-6). **(E-H)** *Vps35*: In male mice, Vps35 gene expression was significantly increased at the ST timepoint (**E.** *p=0.01 with Welch's correction, n=4-6) but was unchanged in LT shRit2 male mice (**F.** p=0.56, n=6-8). In females, Vps35 was unchanged in ST shRit2 female mice (**G.** p=0.81, n=6-7), and was significantly increased by the LT timepoint (**H.** *p=0.04, n=). **(I-L)** *Syt1*: Syt1 gene expression was unchanged in ST shRit2 males (**I.** p=0.46, n=4-6), LT shRit2 males (**J.** p=0.12, n=6-8) or ST shRit2 females (**K.** p=0.93, n=6-8), but was significantly decreased in LT shRit2 females (**L.** **p=0.002, n=5-6).

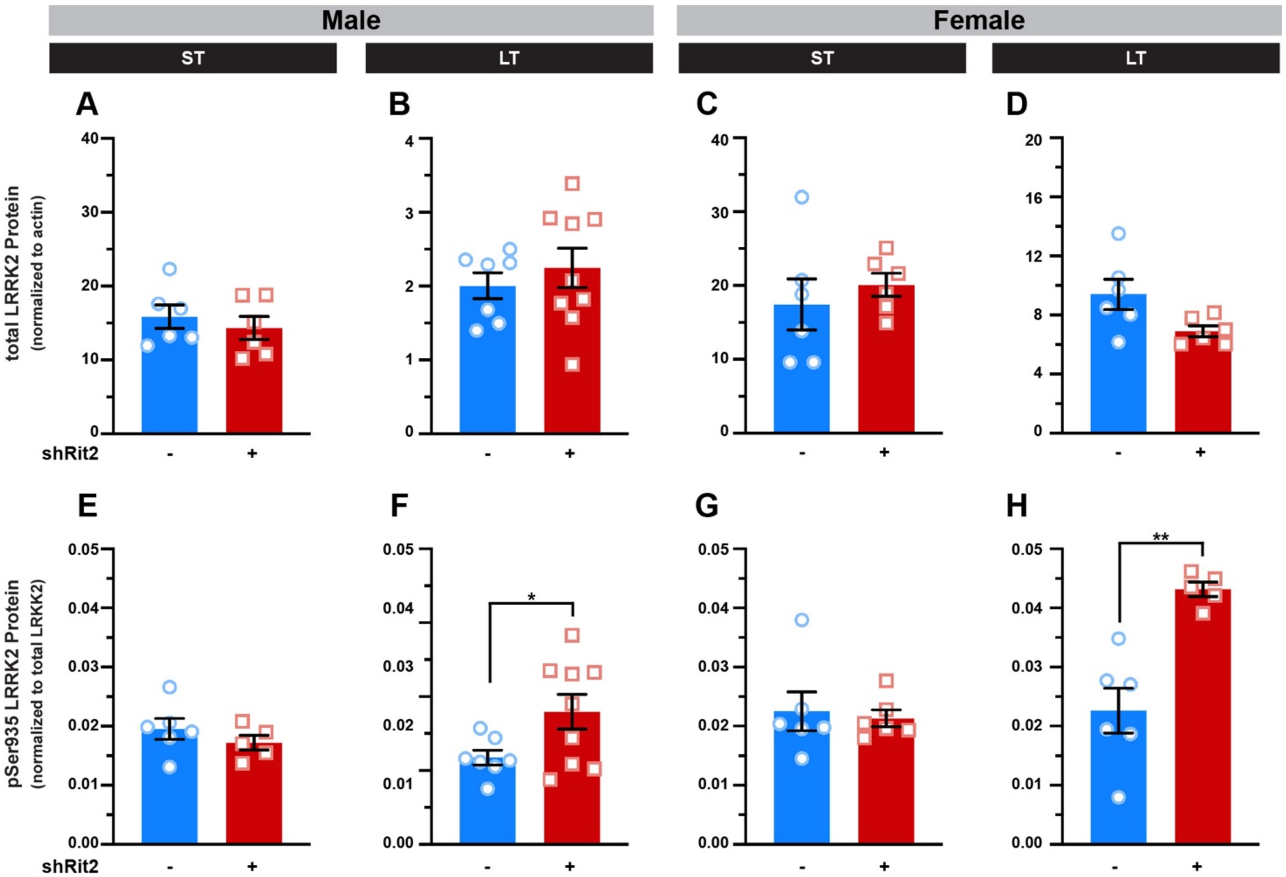


**Supplemental Figure 6. Effect of DAergic Rit2 KD on total and pSer935 LRRK2 protein levels in striatum.** Male and female *Pitx3*$^{IRES\text{-}tTA}$ mouse VTA were bilaterally injected with either AAV9-TRE-eGFP or AAV9-TRE-shRit2 and striatum were dissected from male and female control and shRit2 mice at the indicated timepoints, lysed, and total and pSer935-LRRK2 levels were assessed by quantitative immunoblot as described in *Methods*. Data were analyzed by unpaired, two-tailed Student's t test. N values indicate independent animals **(A-D)** *Total LRRK2 normalized to actin.* Total LRRK2 levels were unaffected by DAergic Rit2 KD in males and females, at either ST or LT timepoints. (**E-H**) *pSer935LRRK2 normalized to total LRRK2.* pSer935LRRK2 levels were normalized to total LRRK2 from (A-D). ST DAergic Rit2 KD had no significant effect on pSer935-LRRK2 levels in either males (**E**) or females (**G**). LT DAergic Rit2 KD significantly increased pSer935-LRRK2 levels in both males (**F;** p=0.04 with Welch's correction, n=7-9) and females (**H**; p=0.002 with Welch's correction, n=5-6).
